# Life on the edge: a new toolbox for population-level climate change vulnerability assessments

**DOI:** 10.1101/2023.06.23.543988

**Authors:** Christopher D. Barratt, Renske E. Onstein, Malin L. Pinsky, Sebastian Steinfartz, Hjalmar S. Kühl, Brenna R. Forester, Orly Razgour

## Abstract

1. Global change is impacting biodiversity across all habitats on earth. New selection pressures from changing climatic conditions and other anthropogenic activities are creating heterogeneous ecological and evolutionary responses across many species’ geographic ranges. Yet we currently lack standardised and reproducible tools to effectively predict the resulting patterns in species vulnerability to declines or range changes.
2. We developed an informatic toolbox that integrates ecological, environmental and genomic data and analyses (environmental dissimilarity, species distribution models, landscape connectivity, neutral and adaptive genetic diversity and genotype-environment associations) to estimate population vulnerability. In our toolbox, functions and data structures are coded in a standardised way so that it is applicable to any species or geographic region where appropriate data are available, for example individual or population sampling and genomic datasets (e.g. RAD-seq, ddRAD-seq, whole genome sequencing data) representing environmental variation across the species geographic range.
3. We apply our toolbox to a georeferenced genomic dataset for the East African spiny reed frog (*Afrixalus fornasini*) to predict population vulnerability, as well as demonstrating that range loss projections based on adaptive variation can be accurately reproduced using data for two European bat species (*Myotis escalerai*, and *M. crypticus*).
4. Our framework sets the stage for large scale, multi-species genomic datasets to be leveraged in a novel climate change vulnerability framework to quantify intraspecific differences in genetic diversity, local adaptation, range shifts and population vulnerability based on exposure, sensitivity, and range shift potential.

## Introduction

Global climate change is affecting biodiversity in unprecedented ways, compounded by other anthropogenic impacts such as habitat degradation, fragmentation and loss (IPBES, 2019). For example, increased temperatures and frequencies of extreme climatic events are predicted to create new selection pressures by rapidly altering resource availability, exposure to pathogens, and the structure and functioning of trophic networks for many species (Bellard et al. 2012; Hoffmann & Sgrò, 2011; Pinsky et al. 2019). How species respond to these new selection pressures depends not only on the magnitude of change occurring, but also on their adaptive capacity (IPCC, 2014), which is influenced by their genetic diversity and phenotypic plasticity (Merilä & Hendry, 2014, Fox et al. 2019), as well as their ability to move and shift their range (Parmesan, 2006; Pecl et al. 2017). Until recently, accounting for these differences in adaptive capacity was largely ignored in climate change vulnerability assessment approaches. However, highlighting intraspecific populations that are most at risk of local extinction, or identifying those with pre-adapted genotypes that can be sources for assisted gene flow and evolutionary rescue (Bell & Gonzalez, 2009), can greatly improve biodiversity conservation management by safeguarding populations and genetic diversity beneficial for resilience to future global change (Hoban et al. 2021, 2022; Laikre et al. 2010).

During the past decade, the dominant approach to climate change vulnerability assessments was based on forecasts of how species ranges are predicted to change using species distribution models (SDMs) (Elith & Leathwick, 2009; Guisan & Thuiller, 2005, Barbet-Massin et al. 2012; Pacifici et al. 2015; Urban, 2015), in some cases refined using genetic data to build SDMs independently for intraspecific populations that have divergent ecological niches (e.g. Bittencourt-Silva et al. 2017; Collart et al. 2021; Ikeda et al. 2017). However, even when accounting for neutral population structure, a major limitation of these approaches has been that intraspecific local adaptation and differential responses to climate change have been largely ignored, potentially leading to inaccurate predictions of future distributions and misplaced conservation efforts (Hällfors et al. 2016; Foden et al. 2019). Adaptation to local environmental conditions is widespread across the tree of life (Hereford, 2009), and the geographic distribution of adaptive variation likely plays a fundamental role in the ability of populations within species to respond to global change (Capblancq et al. 2020; Exposito-Alonso et al. 2018, 2022, Forester et al. 2022). Assessing SDMs together with local adaptation, neutral genetic diversity and predicted range shifts (e.g. Brennan et al. 2022; McGuire et al. 2016; Parks et al. 2022) is therefore essential to understand geographic differences in adaptive capacity and vulnerability under future global change.

Recent calls were made in the emergent field of climate change genomics (Lancaster et al. 2022) for the integration of genomic data to improve the accuracy of climate change vulnerability assessments (Capblancq et al. 2020; Fitzpatrick & Keller, 2015; Nadeau & Urban, 2019; Pauls et al. 2013; Waldvogel, Feldmeyer, et al. 2020). Conceptual and analytical developments enabling the incorporation of intraspecific adaptations across species ranges (e.g. Aguirre-Liguori et al. 2021; Bay et al. 2018; Forester et al. 2023; Razgour et al. 2018, 2019; Ruegg et al. 2018), and phenotypic plasticity (Benito Garzón et al. 2019) have led to major advances in our ability to assess how adaptive capacity and ultimately population vulnerability varies across species ranges. Despite these recent advances, we lack practical and integrative tools to implement analyses across multiple taxonomic groups and geographic regions (see Pinsky et al. 2022). Due to the high multidisciplinarity and diversity of analyses required for most integrated climate change vulnerability assessments, researchers often tailor their approach to their own study system, without creating standardised data structures and code that can be applied more widely to any system (see Waldvogel, Schreiber, et al. 2020, Johnston et al. 2023).

To address this gap in our ability to predict population vulnerability to global change, we introduce ‘Life on the edge’ (hereafter LotE), an analytical toolbox to integrate ecological (species distributions), environmental (environmental dissimilarity and landscape connectivity) and genomic information (neutral and adaptive diversity) in a novel climate change vulnerability assessment toolbox. Throughout the LotE Toolbox, all data inputs are standardised and all code is open source, generalised and parallelised, so it is applicable across any number of different species from any geographic area for which suitable genomic, ecological and environmental data exist. Our toolbox is based on the concepts introduced in Razgour et al. (2018, 2019) to leverage information obtained from the raw data to estimate ‘Exposure’ (estimated from the magnitude of predicted climate change), ‘Sensitivity’ (estimated from neutral and adaptive genetic diversity), and ‘Range shift potential’ (the estimated potential for future distributional shifts and evolutionary rescue given predicted future climate change). Together, exposure, neutral/adaptive sensitivity and range shift potential are combined for each unique location with georeferenced samples to predict population vulnerability to global change across a species’ range. Population vulnerability is defined as the likelihood of a population to become locally extinct.

## Materials and Methods

### Modelling objective

Our overarching goal was to build an informatic toolbox to predict population vulnerability to global change, integrating and generalizing code to make analyses applicable to any suitable population genomic dataset. To build our toolbox we expanded upon two recently published conceptual and analytical frameworks (Razgour et al. 2018, 2019) for climate change vulnerability assessments. Each of the two frameworks integrates genomic and environmental data to assess climate change vulnerability by incorporating a combination of SDMs, landscape connectivity analyses (using electrical circuit theory) and genetic diversity (neutral and adaptive). We integrate SDMs and environmental dissimilarity (‘Exposure’), landscape connectivity (‘Range shift potential’), standing genetic diversity (‘Neutral sensitivity’) and adaptive genetic diversity (‘Adaptive sensitivity’) to estimate population vulnerability for all unique geographic locations with samples across species ranges. Our highly flexible toolbox makes it possible to ‘plug in’ any species with suitable data so that standardised analyses and comparisons across different taxa and regions can be readily made. The LotE Toolbox therefore establishes a backbone for a generalised framework that aims to stimulate a new wave of data synthesis, increasing reproducibility and standardised reporting (Waldvogel et al. 2020) for the research community in population level and species level climate change vulnerability assessments. For ease of interpretation, Fig. 2 summarises the main inputs, analyses and outputs for the toolbox.

**Fig. 1.**
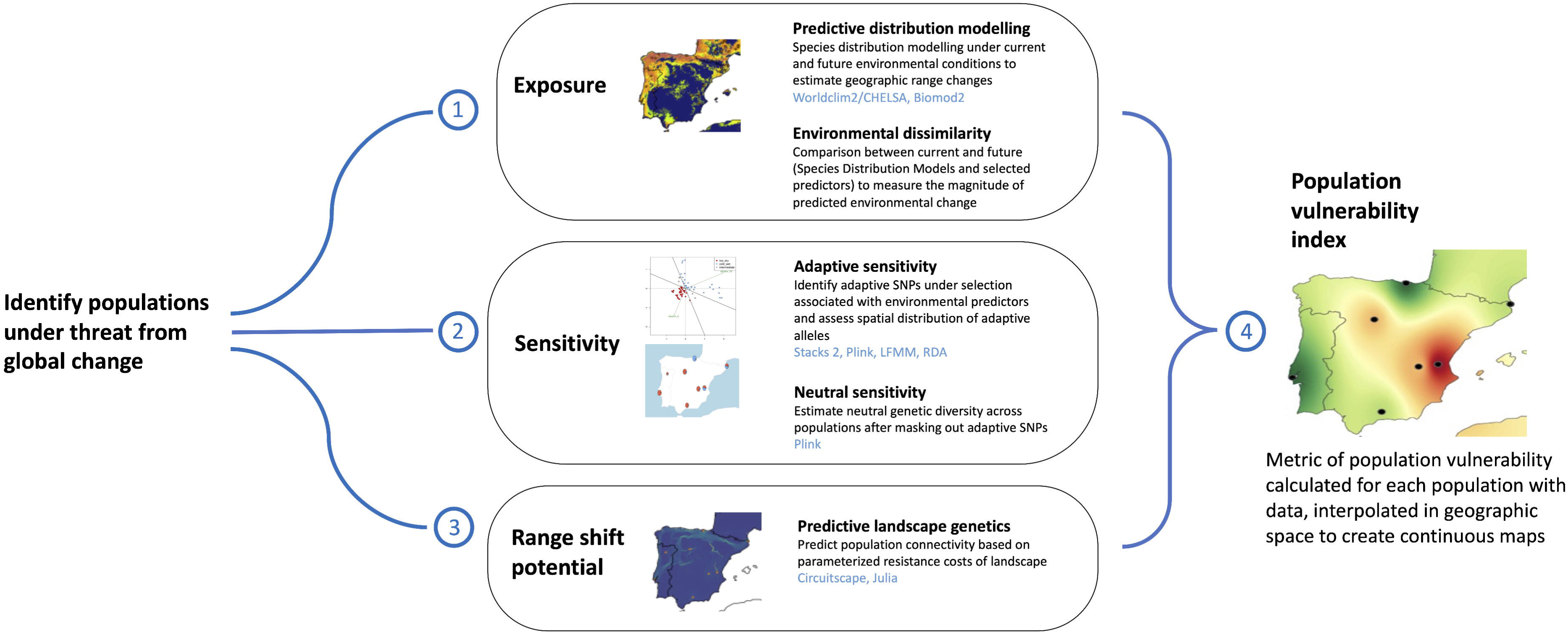
Conceptual and analytical framework for the Life on the edge toolbox, incorporating ‘Exposure’ (current and projected future species distributions and environmental dissimilarity), ‘Sensitivity’ (adaptive alleles and neutral genetic diversity), ‘Range shift potential’ (predicted current and future population connectivity) to predict a final ‘Population vulnerability index’ for each population, interpolated in geographic space. Software packages used are denoted in blue text (LFMM - Latent Factor Mixed Models, RDA - Redundancy Analysis).

**Fig. 2.**
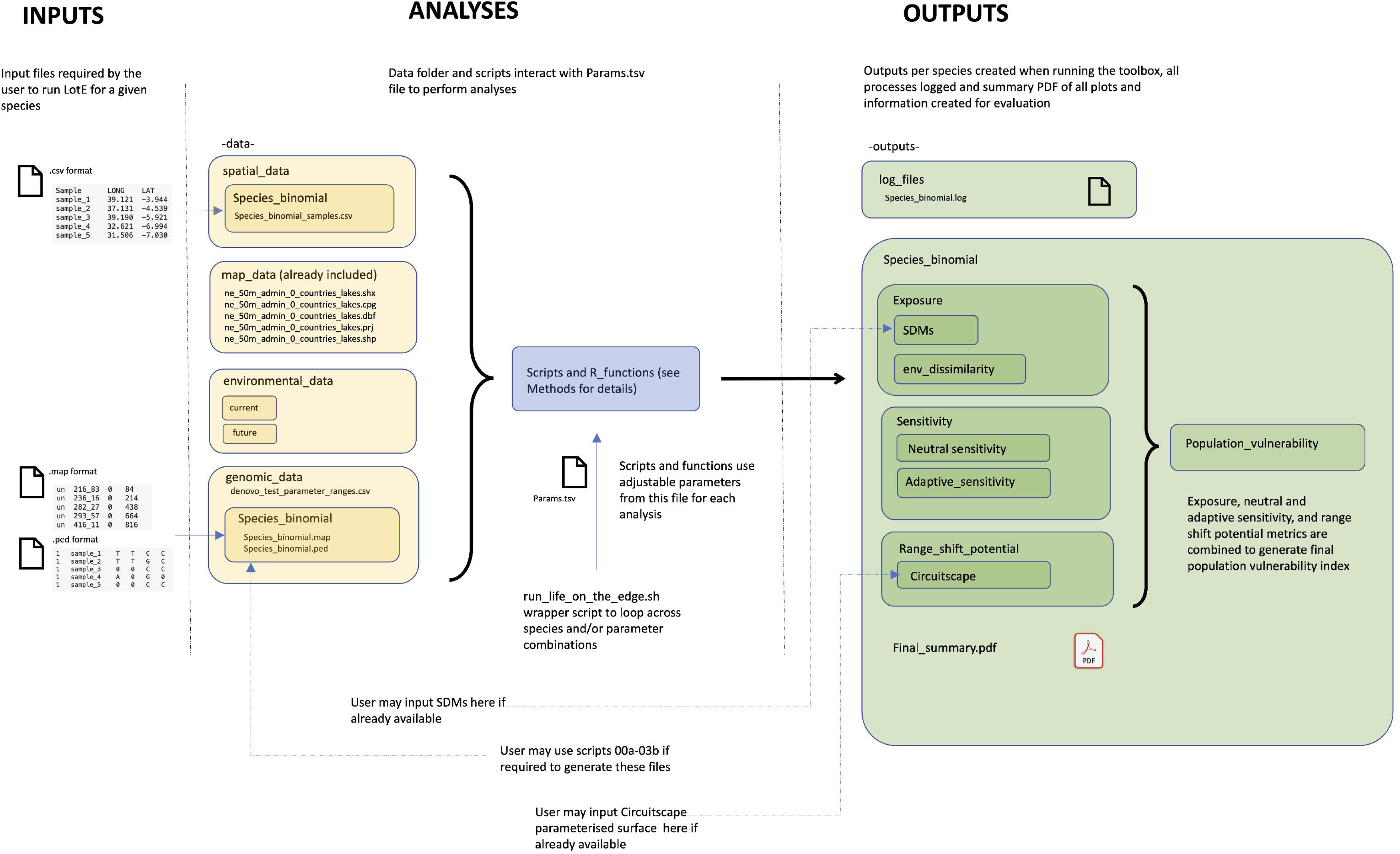
Main inputs and data (yellow boxes), analyses (blue box) and outputs (green boxes) of the LotE Toolbox. Users may store the toolbox in any location, but need to adhere to the directory structure shown above. The *-data-* folder contains four main directories, one of which contains mapping data (world shapefile files available from http://naturalearthdata.com, already provided), another the environmental data, the other two containing folders for each species’ genomic and spatial data. The *-scripts-* and *R_functions* folders contain all the toolbox scripts and functions, and the *-outputs-* folder is created automatically when running the toolbox. Blue lines represent locations for input files, dotted blue lines represent locations for input files if the user wants to circumvent the full toolbox workflow with their own input data.

### Running the toolbox

Life on the edge integrates diverse genomic and spatial analyses to create metrics of exposure, neutral and adaptive sensitivity, range shift potential and population vulnerability. The *run_life_on_the_edge.sh* wrapper script can be used to run each part of the toolbox as required by calling the desired functions. In Supporting Information Text S1 we provide a detailed overview of the input data and structure required and the main methods adopted by the LotE Toolbox at each step, how these are implemented and which aspects are modifiable by the user. Table 1 provides further details on the modelling output, steps taken to achieve that output, and which functions in the toolbox are used at each step. The functions and scripts used (written in R, bash, and Julia) are available in the <u>LotE github website</u> and an accompanying vignette are available at https://cd-barratt.github.io/Life_on_the_edge.github.io/Vignette. The toolbox enables HPC parallelisation, particularly useful for computationally intensive steps. We provide example benchmarking times for analyses to complete (Table 2) to provide users with an overview of processing times.

**Table 1.**
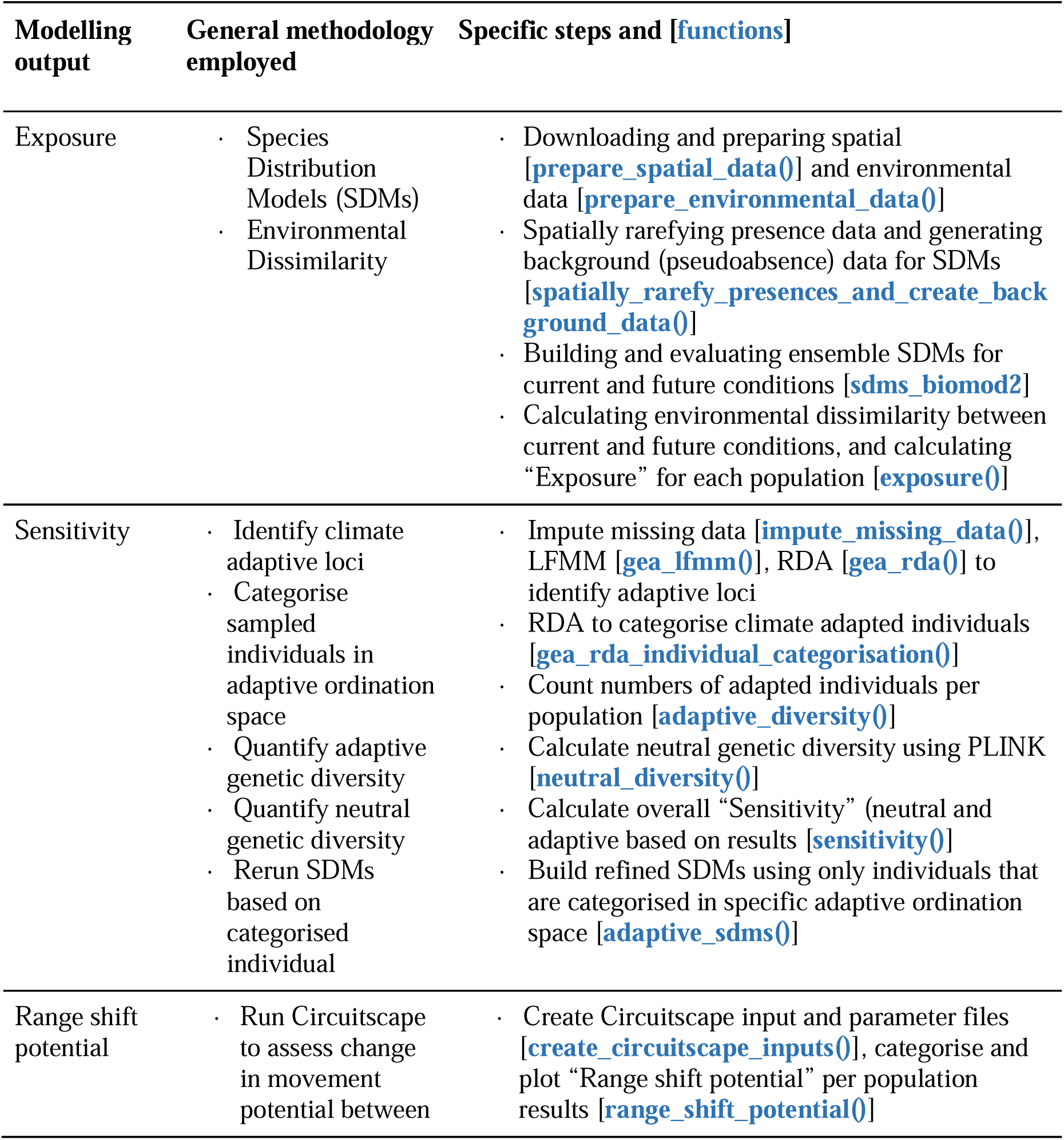
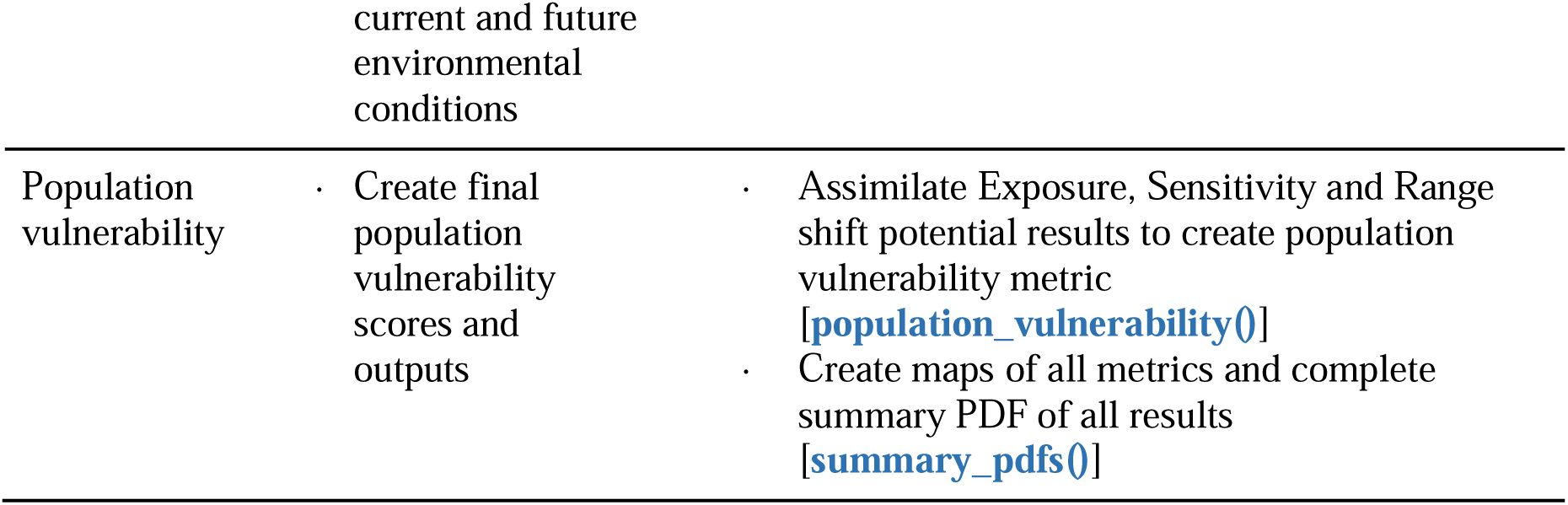
Summary of modelling outputs and methodologies incorporated for the LotE Toolbox. Specific steps for each method are detailed, along with toolbox functions used in each case.

**Table 2.**
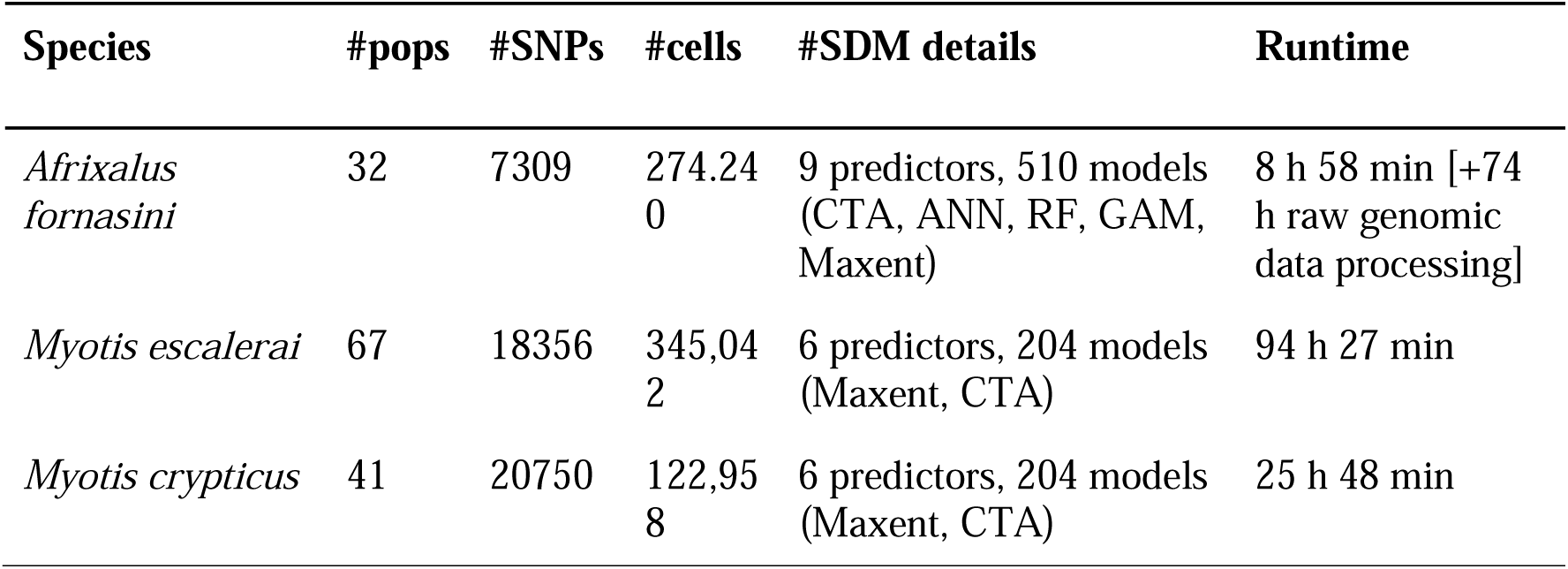
Example benchmarking time for completion of the toolbox on the published data utilised for the three focal species. Genomic data pre-processing times are not included, analyses were each run as separate jobs on a single HPC cluster core (8 threads, NVMe storage) with 80GB RAM allocated.

### Input data

To run the toolbox, appropriate environmental predictor data must first be downloaded (e.g. global data in *.tif* raster format) and stored in the *-spatial_data-/environmental_data* folder, the *params* (Params.tsv) file must be populated with the relevant parameters for each step of the analyses (see Table S1 for a description of each parameter, and the vignette). The three input files required to run a species dataset are the spatial data of the genomic samples (Sample name, Longitude, Latitude, in *.csv* format), and two standard PLINK (Purcell et al. 2007) formatted files *(.ped* and *.map*) for the genomic data (Fig. 2). Details of data naming conventions can be found in the vignette, along with example input data available in the LotE github repository (https://github.com/cd-barratt/life_on_the_edge). To provide reliable outputs using LotE we advise that the genomic samples cover an adequate range of environmental conditions that the species as a whole experiences (i.e. a representative proportion of the species range), and the number of SNPs should be in the order of at least several thousand to detect sufficient signals of local adaptation.

For environmental data, we provide global terrestrial Worldclim2 data (30 arc seconds, clipped to East Africa) for testing purposes. Many users will want different spatial extents or even higher resolution data if available for building accurate SDMs and detecting fine-scale environmental variation and local adaptation across populations on land. Marine or freshwater data could also be used here if users focal taxa are non-terrestrial. These data can be, for example, georeferenced *.tif* files (e.g. bioclim or elevation) downloaded from public databases (e.g. Worldclim2 or CHELSA (Fick & Hijmans, 2017; Karger et al. 2017) for current and selected future conditions and SSP scenarios), or any other predictor data relevant to the study species that is available in raster format (e.g. land cover data). The user should decide on which spatial resolution, future time period and scenario is needed. An R script (**00_process_environmental_data.R**) is provided to assist the user in setting up the environmental predictor data in the correct format (i.e. separating a multi-band file into individual predictors represented by *.tif* files for current and future conditions.

If genomic data needs to be processed (i.e. the user does not yet have *map* and *ped* files with called SNP genotypes for the samples), we provide several scripts for downloading and processing the data. For data downloading (e.g. RAD-seq/ddRAD-seq) the **00a_download_ENA_data.sh** and **00b_download_SRA_data.sh** may be used. In each of these scripts, example data downloads are shown, and the user needs to modify these scripts to target datasets of interest. For European Nucleotide Archive (ENA) downloads, a simple FTP web link to the required data needs to be added. For NCBI Sequence Read Archive (SRA) downloads, the SRA toolkit needs to be first installed, and the SRA run selector may be used in a browser on your local computer to obtain a list of the relevant samples (SraRunTable.txt) belonging to the project, which will then be downloaded. For data processing, **01_process_radtags.sh** (using Stacks) demultiplexes the data from sequenced libraries into files for each sequenced individual, requiring additional barcode information for each individual. After demultiplexing, **02a_denovo_map_test_parameters.sh** runs denovo_map.pl from Stacks multiple times to find optimal values of Stacks core parameters for each dataset. As the optimal parameters to use for Stacks vary across each dataset due to inherent differences in sampling strategy, sequencing effort and natural levels of polymorphism in the sampled populations, it is recommended to explore the dataset using a reduced number of samples before running the full denovo_map.pl script to optimise the SNP datasets for downstream analyses. Following Paris et al. (2018) we focus on the m, M, n and r parameters in Stacks as the focus to explore the data before generating the final SNP datasets for downstream analysis. **02b_extract_results.sh** uses the *stacks-dist-extract* command of Stacks to extract the total numbers of sites, the total number of loci, and the levels of polymorphism in each of these across all of the test runs of denovo_map.pl that were run in 02a_denovo_map_test_parameters.sh. It uses the *Stacks_parameter ranges*.*csv* file with the parameter ranges to test to loop over all parameter combinations, and cleanly summarises the data across each of the test runs into summary files that can be read by the following scripts. **02c_plot_results.R** plots visualisations of the outputs from the previous two scripts. As above, the script reads in the *Stacks_parameter ranges*.*csv* file to set data structures and plot limits, and then uses the generated summary files to create informative plots on how the varying parameters (m, M, n, r) result in different numbers of assembled sites, number of polymorphic sites, % polymorphic loci, total number of SNPs and number of new polymorphic sites iterating across values). The user should visually inspect these plots to establish a parameter set that maximises the amount of useful biological information from the dataset (following Paris et al. 2018). **03a_denovo_map_full.sh** runs denovo_map.pl from Stacks to generate output files (.*ped* and *.map* PLINK format files for downstream analyses) with optimised settings for the dataset using all the samples barcode file. Several bash variables can be defined to specify read/write paths and optimised values of m, M, n and r are set based on the tips mentioned above and the previous plots. **03b_vcf_filtering.sh** enables some post-Stacks filtering using vcftools to further improve the biological interpretability of the data. The script will summarise the levels of missing data across individual samples, enabling the identification and removal of ‘bad apples’ (i.e. samples with a substantial proportion of missing data that may be adversely causing high levels of allelic dropout and lower numbers of loci and SNPs (Cerca et al. 2021).

### Judgement calls to be made by the user

Several important decisions relevant to each species dataset are required throughout the LotE toolbox. Firstly, regarding environmental data, the selection of environmental predictors for SDMs and GEAs should be ecologically relevant to the study species. Predictor variables should be at a suitable spatial grain to be able to detect signals of local adaptation, sufficiently variable between sampled populations, and expected to influence the distribution and/or genetic diversity of the species in question. Secondly, spatial occurrence data (presences) should be checked thoroughly to ensure that incorrect or unrealistic presence data is not included for SDMs (i.e. outside of the native range, or inverted coordinates for example), and that correct taxonomy is followed (e.g. only confirmed species records are included). We have taken measures using the CoordinateCleaner R package (Zizka et al. 2019) to deal with these potential problems, but data should be carefully inspected before analysis and interpretation of results. Additionally, SDMs require consideration of the geographic modelling extent, selection of background (pseudoabsence) data, data partitioning (training vs. testing) strategy and model evaluation in order to follow best practices in the field (see Merow et al. 2013, Araujo et al. 2019, Zurell et al. 2020 for guidelines), and the SDM output itself should be inspected to confirm that it is a reasonable prediction for the species and thus suitable for further use. Thirdly, regarding genomic data, if processing raw sequence read data to generate *.ped* and *.map* files to input for LotE, decisions need to be made to optimise core parameters in Stacks for generating final output files. We advise following our genomic data preparation scripts (00a-03b) which adopt best practices of Paris et al. (2017) and Cerca et al. (2021) to maximise polymorphism in input data while reducing potential ‘false’ loci caused by over- or under merging SNP loci as well as excluding problematic and poorly sequenced samples. For assessing neutral and adaptive sensitivity, including the imputation of missing data and accounting for neutral population structure for GEA analysis, decisions are required to test a reasonable number of genetic clusters (k) represented by the data (i.e. > 1 but less than 10 for example). In the GEA analyses themselves, the thresholds for defining putatively adaptive SNPs are also flexible to enable decisions on how tolerant the user is of False Discovery Rates, defined for each species dataset using the GIF and SD parameters (see Francois et al. 2016, Forester et al. 2018). Finally, parameterisation of landscape resistance surfaces should be based on the ecology of the species in question, with higher resistance values assigned to less permeable landscape/environmental features. The default function for this within LotE is coded to generate resistance surfaces based on the current SDM output, current climate (bioclim 1 and bioclim 12, as well as the selected variables used in the GEA analyses), slope, and land cover which is reclassified based on the ecology of the species (by default less resistance for forest habitats). The resistance surfaces and how they are weighted together to create a cumulative resistance surface for quantifying range shift potential may be parameterised in the *params* file, or alternatively prepared outside the LotE toolbox using software such as ResistanceGA (Peterman, 2018).

### Installation and dependencies

Users of the LotE Toolbox should be proficient in R and bash. As the toolbox utilises several programming languages which in turn require dependencies, initial setup is essential. A working installation of PLINK (Purcell 2007) and Circuitscape (Anantharaman et al. 2019) is required as well as a recent version of R (4.1.3 or later), a bash shell, and a version of Singularity (Kurtzer et al. 2017). Installation of package dependencies from within R needs to be performed upon first running the toolbox (see 00_setup.R). The LotE Toolbox is designed to run in an high performance computing (HPC) environment given the computational resources required especially for large datasets with high numbers of samples (e.g. >250) and sampling localities (>50). For smooth HPC integration we recommend using the supplied Singularity container in the LotE github repository containing a working R version (4.1.3) where relevant R packages are installed and LotE can be run. If required, genomic data processing can be performed in Stacks (Rochette et al. 2019), a bioinformatic pipeline for processing short read (RAD-seq/ddRAD-seq) type data. Many alternative options are available for data processing (e.g. Eaton & Overcast, 2020; McKenna et al. 2010;) which may also cater to non-diploid organisms (e.g. Puritz et al. 2014, Clark et al. 2019) though we focus on Stacks for this toolbox to align with previous analyses (in Razgour et al. 2018, 2019), its wide adoption in the climate change genomics community, and its exceptional documentation and flexibility.

### Modularity

The LotE Toolbox is fully transparent and parameterisable, with standardised workflows following best practices for the processing of genomic data, (Paris et al. 2017, Cerca et al. 2021), species distribution models (Araújo et al. 2019), and genotype-environment association analyses, (Forester et al. 2018, Capblancq & Forester, 2021). Transparency allows all results to be traced back to the data and helps avoid the toolbox being a ‘black box’. We recommend user feedback and sanity checks at several decision-making points of the workflow in order to follow best practices in the relevant subfields of ecology and evolution. Our toolbox can be used as a single pipeline (e.g. from raw sequence and spatial data through to predicting population vulnerability) or in a modular fashion using specific functions. Furthermore, the toolbox offers flexibility, so if users wish to supply their own data (e.g. pre-processed genomic data inputs, their own species presence records, distribution projections, or their own environmental data) it is possible to circumvent steps within the toolbox by simply adding the relevant files to the appropriate directories (see vignette).

### Empirical datasets to demonstrate the utility of the LotE Toolbox

#### Afrixalus fornasini – ‘Novel’ LotE analysis (including genomic data processing)

To demonstrate the utility of the LotE Toolbox we ran the toolbox in its entirety for the East African spiny reed frog, *Afrixalus fornasini*. We processed georeferenced genome-wide RAD-seq data from Barratt et al. (2018) (SRA accession number: PRJNA472166, Table S2) in Stacks, and collated spatial data including published data in Barratt et al. (2018) and cleaned records from the Global Biodiversity Information Facility (GBIF 2020). Environmental data from Worldclim2 was used - bioclim layers 1-19 and slope (Fick & Hijmans, 2017) and land cover (Schipper et al. 2020) at 30s spatial resolution (recategorised into 9 classes following Razgour et al. 2019 to reduce complexity in the model (‘landcov1’) see Table S3). We defined our modelling extent to capture the known range of the species across East Africa, encompassing sampled populations across Tanzania, Kenya, Mozambique and Malawi, used variance inflation factors (VIFs) to reduce spatial autocorrelation in input SDM predictor variables, and assessed local adaptation to maximum temperatures of the warmest month (bioclim_5) and precipitation of the warmest quarter (bioclim_18) as these are known to be important predictors of the species distribution (Barratt et al. 2017, 2018). We parameterised a cumulative resistance surface using five ecologically relevant variables (current SDM, slope, land cover, bioclim_5 and bioclim_18), with respective weights of 0.25,0.1,0.25,0.2,0.2. Exposure, Neutral sensitivity, Adaptive sensitivity, Range shift potential and Population vulnerability were all quantified using the ‘interval’ option, reading defined thresholds for each variable from the *params* file to determine scores. Full parameter settings for the *Afrixalus fornasini* analyses can be found in Table S4.

#### Myotis escalerai and Myotis crypticus – ‘Partial’ LotE analysis - Local adaptation and adaptive SDMs

Secondly, we conducted a ‘partial’ LotE Toolbox run using data from Razgour et al. (2019) (European Nucleotide Archive accession no. PRJEB29086, Table S2) for two European bat species, *Myotis escalerai* and *M. crypticus*. We collated spatial (including cleaned records from the Global Biodiversity Information Facility, GBIF, 2020) and Worldclim2 environmental data at 30s spatial resolution (bioclim_1, bioclim_4, bioclim_7, bioclim_5, bioclim_6, slope, Fick and Hijmans, 2017), as well as land cover (Schipper et al. 2020) (recategorised into 9 classes following Razgour et al. 2019, Table S3). We generated background points and built SDMs using biomod2 (Thuiller et al. 2021), setting SDM parameters to match those used in the original manuscript, namely the spatial modelling extent, predictor variables, future time projections and General Circulation Models, and SDM modelling algorithms as well as the evaluation criteria for SDMs (ROC>0.8). Processed genomic data (*.ped* and .*map* files) from Razgour et al. (2019) were input for the GEA analyses (LFMM and RDA) to investigate local adaptation to hot-dry and cold-wet conditions based on maximum temperatures of the warmest month (bioclim_5) and precipitation of the warmest quarter (bioclim_18). Adaptive SDMs were built using the adaptive_sdms() function of the LotE Toolbox, using individuals parsed into ‘hot-dry’, ‘cold-wet’ adaptive categories for present and future conditions, again following Razgour et al. (2019). Full parameter settings for the *Myotis escalerai and M. crypticus* analyses can be found in Table S4.

## Results

Below we provide details on results for each of the main analyses across the three species datasets we analysed. Results obtained using the LotE Toolbox on empirical data here closely matched those of the published data for *Afrixalus fornasini*, *Myotis escalerai,* and *M. crypticus*, demonstrating that our toolbox is robust as well as being able to integrate diverse analyses to assist predictions of population vulnerability to global change.

### Afrixalus fornasini – ‘Novel’ LotE analysis (including genomic data processing)

After parameter optimization in Stacks (Fig. S1A) we opted to select Stacks core parameters of M=2, m=5 and r=80% for downstream analyses due to the trade-off between maximizing polymorphism in our data and reducing potential ‘false’ loci caused by over- or undermerging loci. Our final filtered dataset contained 7,309 loci for 43 individuals, and we retained the first SNP from each locus in our final SNP dataset to maintain assumptions of linkage disequilibrium.

Future forecasts of the SDMs showed the majority of the core *Afrixalus fornasini* distribution will remain largely unchanged from current conditions, with some range loss towards central Tanzania (Fig. 3A), and expansions throughout northern Mozambique, parts of coastal Tanzania and southern Kenya (Fig. 3A). Most populations will experience similar warming and drying in the future compared to current conditions, with the exception of populations towards central Tanzania (also within the range contraction areas in the SDMs) and northern Mozambique, which are projected to experience more warming and rainfall loss than the remaining populations (Fig. 3A). Based on our GEA analyses, populations in Northern Tanzania and Kenya are not strongly locally-adapted to climate, falling within the ‘intermediate’ category (Fig, 3B). Hot-dry adapted populations were identified across most coastal Tanzanian populations, whereas colder and wetter adapted populations were located in more mountainous regions (Udzungwa and Uluguru mountains and surrounding areas in Tanzania, and throughout northern Mozambique (Mount. Mabu, Nampula) and southern Malawi (Thyolo, adjacent to Mt. Mulanje). Range shift potential analyses showed generally high connectivity between populations (Fig. 3C), which makes sense ecologically given that *A. fornasini* is a cosmopolitan species that is found in gradients of forest, savannah and bushland habitats (IUCN, 2018, Barratt et al. 2018) and can therefore deal with a high degree of habitat modification and fragmentation. Most populations demonstrated high landscape connectivity with the exception of populations in coastal Tanzania (Lindi, Kilwam Dar, Bagamoyo) or in montane regions of Tanzania (Udzungwa, Nguru, Nguu) which were more isolated from their conspecific populations.

**Fig. 3.**
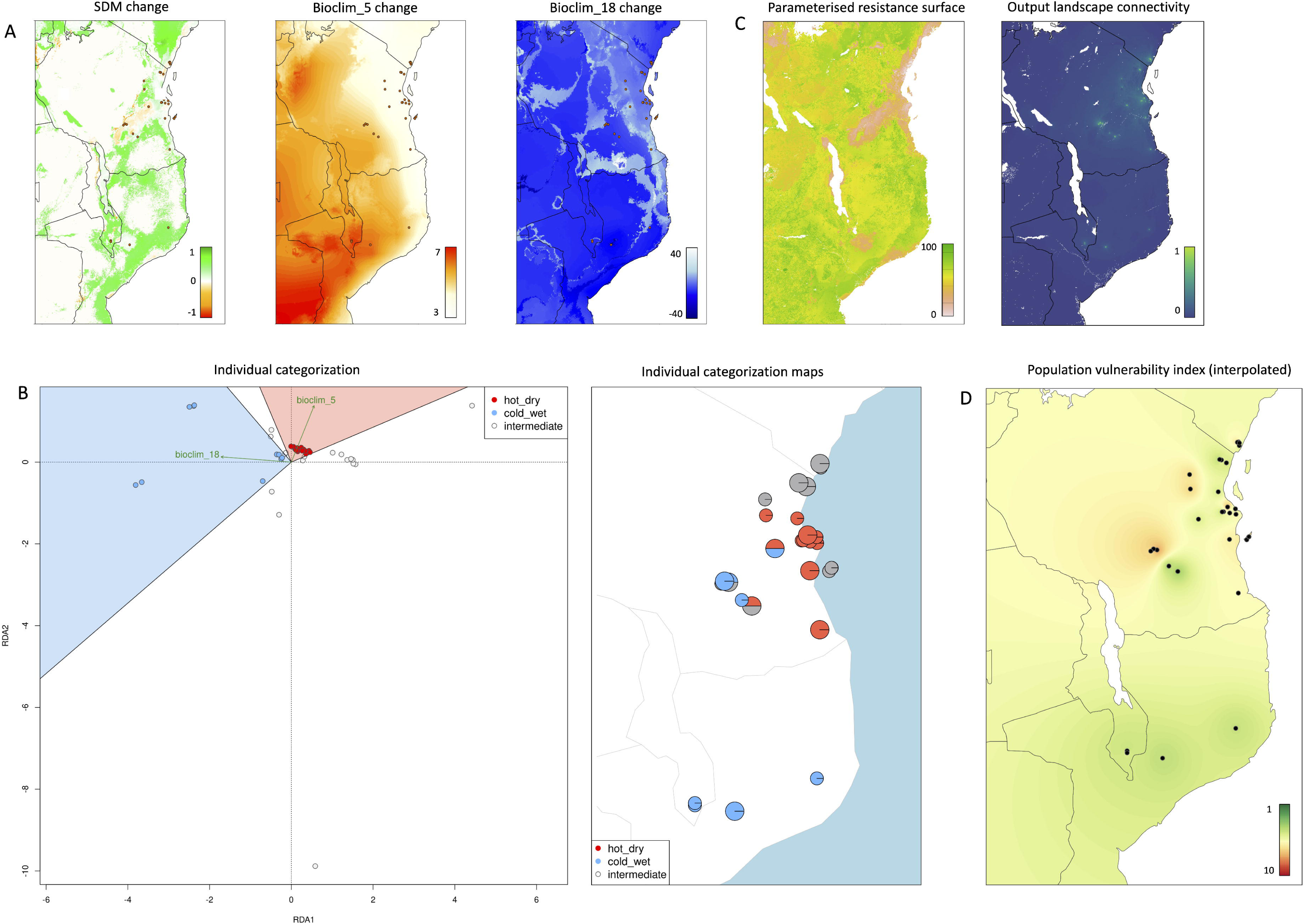
*Afrixalus fornasini* results generated using the LotE Toolbox. A) Exposure - environmental dissimilarity between current and future conditions (SDM (−1 = range loss, 1 = range expansion), maximum temperature of the warmest month (Bioclim_5, °C), precipitation of the warmest quarter (Bioclim_18, mm)). B) Sensitivity – adaptive categorisation of individuals into cold-wet, intermediate and hot-dry genotypes and their distribution across geographic space. C) Range shift potential – parameterised cumulative resistance surface (ranging from 0 - no resistance, to 100 - complete barrier) and predicted movement density between populations based on Circuitscape analysis. D) Population vulnerability, calculated as the mean of the exposure, adaptive and neutral sensitivity, and range shift potential metrics (all ranging between 1 (low vulnerability) and 10 (high vulnerability). Sampling locations with genomic data represented over maps as dots, legends within each panel and plot provide information on the scale of variables.

Our summary maps of exposure, neutral and adaptive sensitivity, and range shift potential based on the sampled populations (Fig. S1C) show that projected future climate change will play a strong role in increasing the exposure of populations in central Tanzania, southern Malawi and northern Mozambique. This extends to reducing range shift potential (i.e. landscape connectivity) for these populations and also for many additional coastal Tanzanian populations. Neutral genetic diversity is mixed across the populations, with lower sensitivity (i.e. higher heterozygosity) in higher elevation regions (e.g Udzungwa, Nguu, Usambara mountains, Muyuyu and Ruvu forest reserves in Tanzania, Thyolo and Nampula in Mozambique) and higher neutral sensitivity scattered throughout the remainder of the coastal forests of Tanzania and southern Kenya (Fig. S1C). Given the relatively low degree of local adaptation, most populations have low adaptive sensitivity except for Mafia island and Tanga region (Tanzania). Together, population vulnerability indices (Fig. 3D) for *A. fornasini* demonstrate that the most at-risk populations given predicted future global change are at the periphery of the species current range (Thyolo in Malawi, Mt. Mabu in Mozambique, Kilombero valley and lowland Usambara mountains in Tanzania). These populations generally demonstrate higher exposure to future climates relative to current conditions, and/or have lower standing genetic diversity (heterozygosity), as well as lower levels of range shift potential compared to other conspecific populations. Full log file output from the ‘novel’ LotE analysis for *A. fornasini* (Appendix S1) as well as a final summary PDF (Appendix S2) can be found in the Supporting Information.

### Myotis escalerai and Myotis crypticus – ‘Partial’ LotE analysis - Local adaptation and adaptive SDMs

After inspecting p-value distributions and adjusting the genomic inflation factor and standard deviation (SD±3) to control false discovery rate (FDR<0.05) thresholds and candidate SNP detection for both local adaptation methods, we detected 79 RDA and 385 LFMM SNPs and 104 RDA and 176 LFMM SNPs for *Myotis escalerai* and *M. crypticus*, respectively. Using a conservative approach (i.e. retaining only loci that were detected across both methods, n=50 and n=26), we parsed individuals into the broad adaptive categories reported in Razgour et al. (2019) (Fig. 4). Of these, 60 *M. escalerai* and 10 *M. crypticus* individuals were adapted to ‘hot-dry’ conditions, 108 *M. escalerai* and 14 *M. crypticus* were ‘cold-wet’ adapted, and 48 *M. escalerai* and 26 *M. crypticus* were categorised as ‘intermediate*’* (Fig. 4A,C). Mapping these individuals in geographic space showed high concentrations of local adaptation to hot-dry conditions in southern and western sampling, and local adaptation to cold-wet conditions in northern and eastern sampling for *M. escalerai*, and for *M*. *crypticus*, hot-dry individuals in northern Spain, cold-wet individuals towards the Pyrenees, closely matching the results from Razgour et al. (2019) (Fig. 4B,D).

**Fig. 4.**
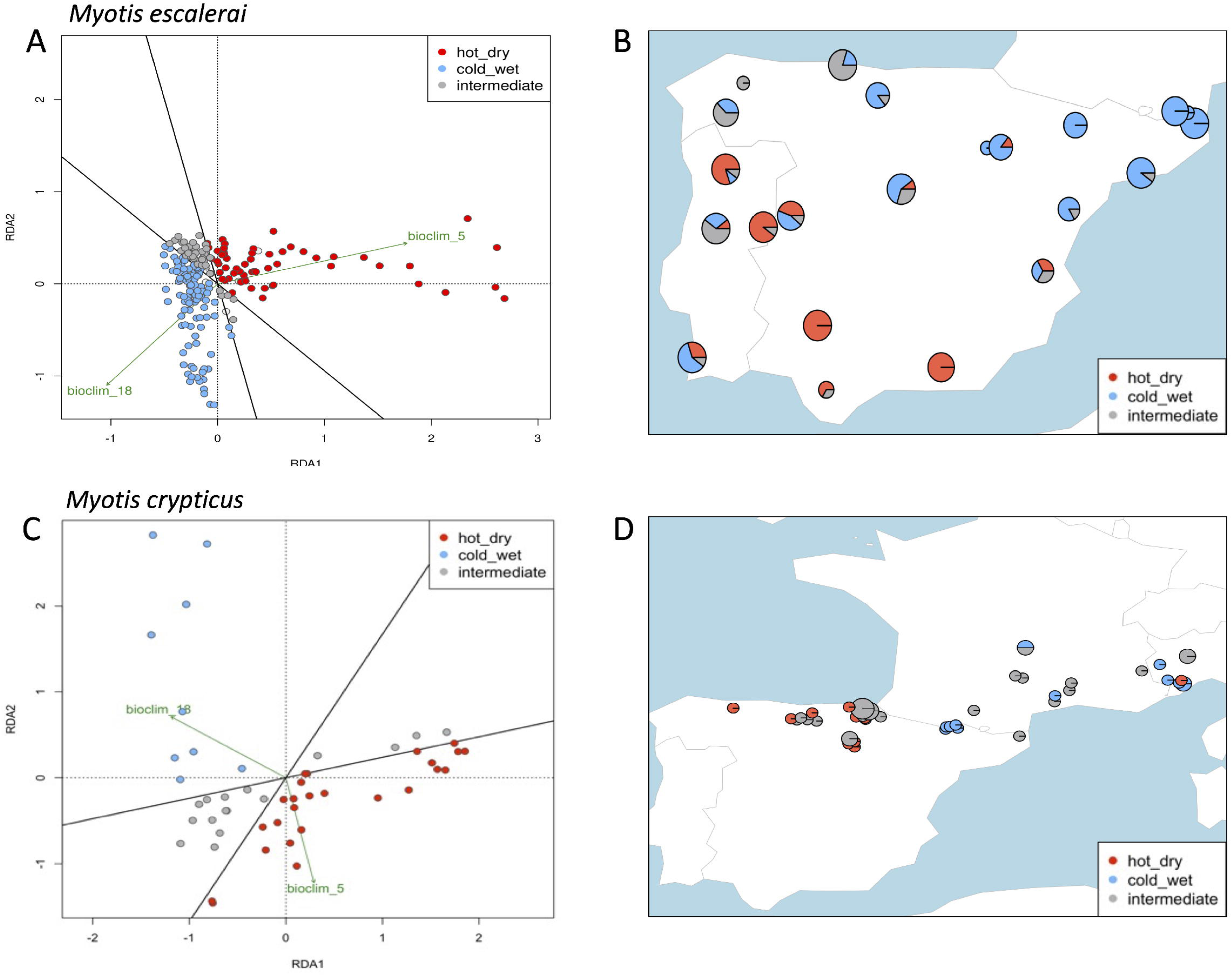
Individual categorization results for *Myotis escalerai* and *Myotis crypticus* generated using the LotE Toolbox. A) *M. escalerai* individual categorisation in RDA ordination space based on putatively adaptive SNPs and B) Mapped categorized individuals in geographic space. C) *M. crypticus* individual categorisation in RDA ordination space based on putatively adaptive SNPs and D) Mapped categorized individuals in geographic space.

Species distribution models predicted future contractions of potential habitat suitability for *M. escalerai* across most of its range in the Iberian peninsula, particularly in more arid regions (southern Spain and Portugal), and expansions predicted in northern Portugal and parts of the Pyrenees (Fig. S1A). *M. crypticus* habitat suitability also decreased in future climate projections, with the Iberian region predicted to be largely unsuitable and future suitability limited to elevated regions including parts of the Alps and the Pyrenees. Separating the different categories of individuals (hot-dry and cold-wet adapted), both species results broadly matched Razgour et al. (2019) (Fig. 5), in *M. escalerai,* suitable environmental conditions for both categories were predicted to shift northwards, substantially affecting the predicted ranges, in particular the habitat suitability for cold-wet genotypes found in northern Iberia, which substantially contracted in future conditions (Fig. 5A,B). *M. crypticus* showed high habitat suitability in northern parts of the Iberian peninsula, the Pyrenees, which were predicted to contract in future conditions, and parts of southern Europe, which were predicted to shift northwards in the future (Fig. S1B). As with *M. escalerai,* hot-dry and cold-wet adapted individuals’ habitat suitability was predicted to decrease slightly, particularly for the latter, whose potential suitable range throughout the Pyrenees and central and south-western France almost completely disappeared under future predictions (Fig. 5A,B). Full log file outputs from the ‘partial’ LotE analysis for *M. escalerai* and *M. crypticus* can be found in the Supporting Information (Appendix S3, S4).

**Fig. 5.**
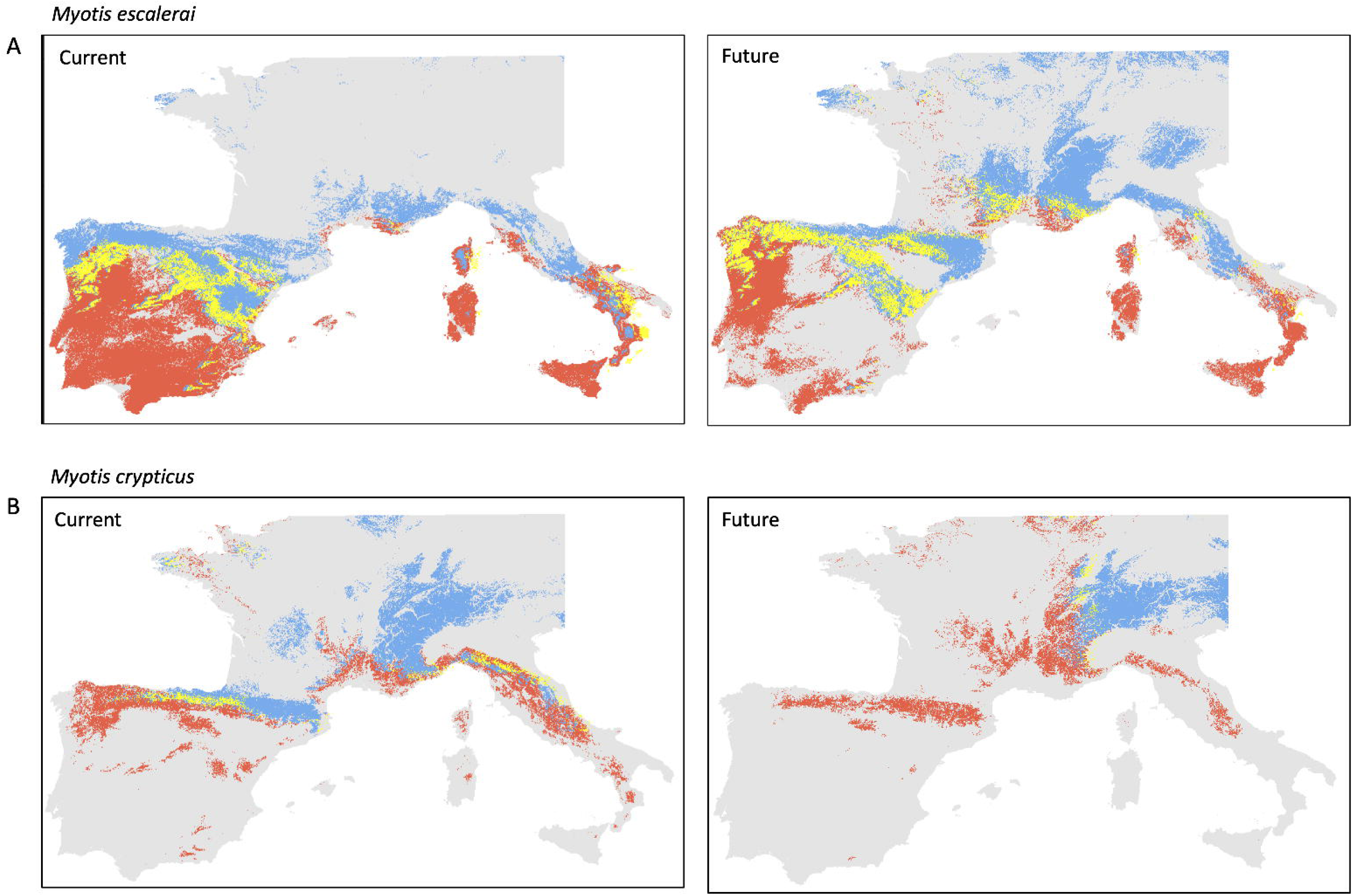
Adaptive SDMs generated using the LotE Toolbox capturing intraspecific adaptations for *Myotis escalerai* and *Myotis crypticus* based on the categorized individuals for hot-dry, cold-wet conditions shown in Fig. 4. Maps are categorized into binary presence/absences for hot-dry adapted (red), cold-wet adapted (blue), with overlapping areas for both categories in yellow. A) *M. escalerai* adaptive SDMs (left panel: current conditions, right panel: future conditions). B) *M. crypticus* adaptive SDMs (left panel: current conditions, right panel: future conditions).

## Discussion

Current climate change genomics approaches to assess population vulnerability lack practical and integrative tools to implement analyses across multiple taxonomic groups and geographic regions (Pinsky et al. 2022). Adaptive responses to global change are likely to be insufficient for many species (Quintero & Wiens, 2013; Radchuk et al. 2019), and we need standardised climate change vulnerability tools incorporating genomics with ecological and environmental data that can be applied across diverse datasets (i.e. species/regions). Previous studies have predicted population vulnerability by developing custom frameworks applied to specific study systems (e.g. Aguirre-Liguori et al. 2021; Bay et al. 2018; Razgour et al. 2018, 2019; Ruegg et al. 2018). With Life on the Edge, we present a novel, generalised and customizable toolbox which can perform sophisticated analyses in a straightforward and reproducible way. The toolbox enables large scale data synthesis across multiple species and geographic areas and the outputs address ecological and evolutionary responses to future global change and can be used to guide conservation action. We envisage LotE as a key tool in the emergent field of climate change genomics, with scope for future development and expansion of the concepts presented in this manuscript. Below, we discuss applications of the LotE Toolbox in its current form, its limitations, and future additions, analytical and conceptual developments.

### Current applications for LotE

Life on the edge, with its geographical and taxonomic flexibility, contributes to answering several fundamental questions in ecology, evolution and conservation. Outputs from LotE may therefore be utilised for informing the management of populations to reduce the accumulation of deleterious alleles (genetic load) and maintain genetic diversity (e.g.heterozygosity) within and between populations (e.g. Segelbacher et al. 2022). For example, suitable donor and recipient populations for translocations/evolutionary rescue may be selected based on a combination of heterozygosity estimates, local adaptations and spatial connectivity. With these strategies, conservation managers may avoid introducing maladapted individuals to unsuitable climatic conditions (Chen et al. 2022a; Chen et al. 2022b), thus strengthening the overall individual fitness, genetic diversity, and adaptive potential of populations (Frankham, 2015; Frankham et al. 2019).

At a regional scale, spatial conservation planning can benefit from the climate change genomics perspective of the outputs generated by the LotE Toolbox. For example, any of the interpolated metrics for neutral and adaptive genetic sensitivity or population vulnerability are useful for prioritising areas that support particularly high levels of genetic diversity or vulnerable populations, using common software such as Zonation, Marxan and relatives (Lehtomäki & Moilanen, 2013; Moilanen et al. 2005, Ball et al. 2009). When data from multiple species in an ecological community are available, evaluating congruence in metrics across species would provide a representative measure of community-level genetic diversity and vulnerability based on genome-wide data (e.g. Schielzeth & Wolf, 2021; Stange et al. 2021). This could be scaled up (i.e. across countries and continents) with sufficient data across taxonomic groups and can be used to identify hotspots of adaptive potential versus regions under threat. Similarly, investigating the ecological, environmental and anthropogenic correlates of LotE outputs can identify causal drivers of observed vulnerability patterns for species and ecological communities (similar to approaches in Howard et al. 2020; Maxwell et al. 2016; Tilman et al. 2017), which can inform broad conservation actions as well as understanding species-specific drivers of declines more thoroughly with population level data.

Finally, the adaptive SDMs within the LotE Toolbox contribute to more realistic estimates of range shifts under global change scenarios, providing improved predictions of future biodiversity losses which may be offset with appropriate conservation measures (Hoffman and Sgro, 2011). Overlooking intraspecific population variability, in particular local adaptation, can result in an overestimation of future biodiversity losses (Razgour et al. 2019), and it is increasingly clear that predictive models informed by empirical genomic data provide a more realistic alternative to simplistic modelling approaches that do not account for local adaptation (Forester et al. 2023, Bay et al. 207).

### Future directions

In addition to applications of the current toolbox, there are three future conceptual and analytical developments that could expand the framework and its long-term impact and benefit to the research and conservation community. The toolbox has been purposely designed to be dynamic, so that additional ‘modules’ may be created and integrated in future versions, thus enabling it to evolve in tandem with the research community and adopt best practices and state-of-the-art tools. New methods or tools (for example a new and improved SDM or GEA package) that supersedes existing methods may be integrated relatively simply into updated versions of the toolbox by creating additional functions to supplement or update the existing modules. This also applies to the type of genomic data that can be analysed, which can be scaled up from short read (RAD-seq) type data to whole genome sequencing data simply by switching the bioinformatic processing tool (e.g. from Stacks to GATK, McKenna et al. 2010, for example).

A primary focus for expanding LotE in the future is for monitoring biodiversity change and population vulnerability over time, for example by linking outputs with global conservation efforts such as the Sustainable Development Goals as well as ‘Essential Biodiversity Variables’ (Hoban et al. 2022). By integrating time series data and multiple biological replicates, it would be possible to track changes in exposure, sensitivity (adaptive/neutral), range shift potential, and population vulnerability as new data become available. Although suitable population-level genomic and spatial sampling time-series replicates are currently rare (but see Pfenniger et al. 2023), ever improving advances in sequencing technologies and the exponential accumulation of these data in online repositories (e.g. NCBI Sequence Read Archive, ENI European Nucleotide Archive, DNA Databank of Japan) make the tantalizing prospect of genomics-informed biodiversity monitoring an achievable target in the near future (Bálint et al. 2018; Taus et al. 2017; Pfenniger and Bálint, 2022). Ultimately, automated ‘scraping’ of public repositories for species datasets as new data become available could provide a near real-time assessment of species’ biodiversity status based on the latest available genomic and spatial information, similar to the explosion of data generation, availability and automation to monitor recent epidemiological outbreaks (e.g. http://nextstrain.org).

Secondly, validating the outputs of LotE using sensitivity analyses and simulations will substantially improve predictions and confidence intervals in results (Hoban, 2014). LotE is amenable to parallelization, enabling it to run across a range of parameter settings in the *params* file, with which results can be harvested and parameter variation effects on results can be explored in detail. The integrative analyses of LotE may complement and enhance similar frameworks investigating genomic offset (e.g. Smith et al. 2021) that do not consider migration and gene flow (e.g. Fitzpatrick & Keller, 2015; Rellstab et al. 2021; Capblancq et al. 2020), and could be evaluated against outputs from comparable frameworks using the same underlying datasets. With respect to input genomic data, we acknowledge there has been significant debate on the power of reduced representation library (RAD/ddRAD-seq) datasets to detect sufficient signals of local adaptation (Lowry et al. 2017, but see Catchen et al. 2017). Ideally, whole genome sequencing data would be the gold standard for detecting local adaptation in populations, and scaling our framework to whole genome sequencing datasets will improve the ability to detect both weak and strong signals of local adaptations. Given that these kinds of datasets for range-wide population sampling are presently rare, reduced representation library datasets currently offer the most feasible approach to synthesise data across taxa and regions. Furthermore, an important step could be to integrate simulation studies using artificial fragmentation of whole genome sequencing datasets combined with power analyses (e.g. Patton et al. 2019) to help understand how signals of local adaptation using reduced representation library data such as those demonstrated here can adequately detect local adaptations and where the drop-off in statistical power lies from whole genome datasets (e.g. Benjelloun et al. 2019).

Thirdly, integrating phenotypic plasticity and functional genomics data are a rich potential avenue for expansion for the LotE Toolbox, for example in model systems where there is adequate knowledge on physiological limits for species, or the underlying genetic basis of their functional traits and reaction norms (e.g. Oomen & Hutchings, 2022). Expansion of the toolbox to incorporate this information, for example isolating genomic regions related to thermal stress tolerance and tracking how these vary and are distributed geographically across populations (e.g. Pimsler et al. 2020), would substantially add to the analytical power of the LotE Toolbox, and provide integration with phenotypic plasticity, functional trait and ecological knowledge for model systems. Furthermore, different kinds of ‘omics data (e.g. structural variants such as copy number variants or epigenetic data) could broaden the approach to investigate other types of genomic variants that influence adaptive responses (Layton & Bradbury, 2022; Wollenberg Valero et al. 2022), especially when available from multiple individuals and populations across a species’ range.

### Main considerations when using the LotE Toolbox

There is no ‘silver bullet’ solution to predicting vulnerability to global change. Each dataset input into LotE or any other climate change vulnerability framework has its own idiosyncrasies and biases, and these should be taken into account when drawing conclusions, ideally using sensitivity analyses to investigate the effects of specific parameters at each step. Though no dataset is perfect, we believe the LotE Toolbox can make the most of available datasets at present, and due to the large number of reduced representation short read library datasets (i.e. RAD/ddRAD-seq) published over the past decades (more than 2400 articles as of the end of 2017, Campbell et al. 2018) to assess population structure, phylogeography, demographic history), these presently provide the most promising and widely applicable datasets for our approach. As sequencing technologies continue to improve, we will undoubtedly move towards larger numbers of whole genome sequencing datasets which will provide higher resolution data for assessing genetic diversity and local adaptation in particular.

## Conclusions

We introduce the ‘LotE’ conceptual and analytical framework, a toolbox which facilitates the integration of environmental, molecular and ecological data to perform genomics-informed climate change vulnerability assessments. Given the sheer number of analyses, data preparation steps and computational power required to perform climate change vulnerability assessments, our HPC-compatible framework is automated and standardised, but flexible, thus making it possible to perform comparisons across species and geographic regions. The modular structure of LotE mean that it is not restricted to being a climate change vulnerability assessment tool, but can also be used for batch preparation and analysis of spatial-environmental data, building species distribution models or Circuitscape analyses, investigating local adaptation, or mapping intraspecific neutral and adaptive genetic diversity across species ranges. With the increasing availability of high-quality georeferenced genome-wide datasets published in open access online repositories, as well as constantly improving climate model simulations, the LotE framework offers a range of tools that can be used to investigate intraspecific responses to global change, thus providing empirical results from large genomic and spatial datasets to inform and assist biodiversity conservation in our rapidly changing world. Many opportunities for integrating simulations, functional genomics data, and biodiversity monitoring are possible in the future, and we envisage that LotE will be a useful tool for both the academic research and conservation practitioner communities that can stimulate a new wave of data synthesis, increasing reproducibility and standardised reporting when assessing intraspecific diversity and vulnerability to global change.

## Acknowledgements

All computation was conducted on the UFZ EVE cluster in Leipzig, Germany. The findings and conclusions in this article are those of the authors and do not necessarily represent the views of the US Fish and Wildlife Service.

## Author contributions

CDB, OR and REO conceived the idea of the study and obtained funding for the LotE project along with SS, HK, BF and MP. CDB and OR supplied datasets for the data curation and creation of the LotE Toolbox, with base code from OR and BF. CDB performed all analyses and wrote the manuscript, with contributions from all co-authors. All authors approved the final manuscript.

## Statement on inclusion

Our study brings together authors from a number of different countries but does not include scientists based in the countries where the samples were collected. The toolbox described in this manuscript is mainly methodological and the underlying data was already published, and we were unable to sufficiently involve local researchers due to scarcity of the relevant coding expertise. However, as well as building on and citing previous work by local scientists, the toolbox in this manuscript is currently being used for region and species-specific questions in those respective countries, which do integrate local scientists prominently.

## Data availability

We supply a DRYAD data repository containing all the input files necessary to duplicate the analyses in this manuscript as well as a comprehensive tutorial (https://cd-barratt.github.io/Life_on_the_edge.github.io/). All code is openly available as a Zenodo repository (all repositories are linked at the LotE website). Raw sequence data is available at the European Nucleotide Archive (ENA): *Myotis escalerai* and *M. crypticus* (PRJEB29086), and the NCBI Short Read Archive (SRA): *Afrixalus fornasini –* (SRP150605).

## References

Aguirre-Liguori, J. A., Ramírez-Barahona, S., & Gaut, B. S. (2021). The evolutionary genomics of species’ responses to climate change. Nature Ecology & Evolution, 5(10), Article 10. https://doi.org/10.1038/s41559-021-01526-9

Aiello-Lammens, M. E., Boria, R. A., Radosavljevic, A., Vilela, B., & Anderson, R. P. (2015). spThin: An R package for spatial thinning of species occurrence records for use in ecological niche models. Ecography, 38(5), 541–545. https://doi.org/10.1111/ecog.01132

Anantharaman, R., Hall, K., Shah, V., & Edelman, A. (2019). Circuitscape in Julia: High Performance Connectivity Modelling to Support Conservation Decisions (arXiv:1906.03542). arXiv. https://doi.org/10.48550/arXiv.1906.03542

Araújo, M. B., & New, M. (2007). Ensemble forecasting of species distributions. Trends in Ecology & Evolution, 22(1), 42–47. https://doi.org/10.1016/j.tree.2006.09.010

Araújo, M. B., Anderson, R. P., Márcia Barbosa, A., Beale, C. M., Dormann, C. F., Early, R., Garcia, R. A., Guisan, A., Maiorano, L., Naimi, B., O’Hara, R. B., Zimmermann, N. E., & Rahbek, C. (2019). Standards for distribution models in biodiversity assessments. Science Advances, 5(1), eaat4858. https://doi.org/10.1126/sciadv.aat4858

Bálint, M., Pfenninger, M., Grossart, H.-P., Taberlet, P., Vellend, M., Leibold, M. A., Englund, G., & Bowler, D. (2018). Environmental DNA Time Series in Ecology. Trends in Ecology & Evolution, 33(12), 945–957. https://doi.org/10.1016/j.tree.2018.09.003

Ball, I. R., Possingham, H. P., & Watts, M. E. (2009). Marxan and relatives: Software for spatial conservation prioritization. In A. Moilanen, K. A. Wilson, & H. P. Possingham (Eds.), Spatial conservation prioritisation: Quantitative methods and computational tools (pp. 185–210). Oxford University Press.

Barbet-Massin, M., Jiguet, F., Albert, C. H., & Thuiller, W. (2012). Selecting pseudo-absences for species distribution models: How, where and how many? Methods in Ecology and Evolution, 3(2), 327–338. https://doi.org/10.1111/j.2041-210X.2011.00172.x

Barratt, C. D., Bwong, B. A., Jehle, R., Liedtke, H. C., Nagel, P., Onstein, R. E., Portik, D. M., Streicher, J. W., & Loader, S. P. (2018). Vanishing refuge? Testing the forest refuge hypothesis in coastal East Africa using genome-wide sequence data for seven amphibians. Molecular Ecology, 27(21), 4289–4308. https://doi.org/10.1111/mec.14862

Barratt, C. D., Bwong, B. A., Onstein, R. E., Rosauer, D. F., Menegon, M., Doggart, N., Nagel, P., Kissling, W. D., & Loader, S. P. (2017). Environmental correlates of phylogenetic endemism in amphibians and the conservation of refugia in the Coastal Forests of Eastern Africa. Diversity and Distributions, 23(8), 875–887. https://doi.org/10.1111/ddi.12582

Barratt, C. D., Lester, J. D., Gratton, P., Onstein, R. E., Kalan, A. K., McCarthy, M. S., Bocksberger, G., White, L. C., Vigilant, L., Dieguez, P., Abdulai, B., Aebischer, T., Agbor, A., Assumang, A. K., Bailey, E., Bessone, M., Buys, B., Carvalho, J. S., Chancellor, R., … Kühl, H. (2021). Quantitative estimates of glacial refugia for chimpanzees (Pan troglodytes) since the Last Interglacial (120,000 BP). American Journal of Primatology, 83(10), e23320. https://doi.org/10.1002/ajp.23320

Bay, R. A., Rose, N. H., Logan, C. A., & Palumbi, S. R. (2017). Genomic models predict successful coral adaptation if future ocean warming rates are reduced. Science Advances, 3(11), e1701413. https://doi.org/10.1126/sciadv.1701413

Bay, R. A., Harrigan, R. J., Underwood, V. L., Gibbs, H. L., Smith, T. B., & Ruegg, K. (2018). Genomic signals of selection predict climate-driven population declines in a migratory bird. Science, 359(6371), 83–86. https://doi.org/10.1126/science.aan4380

Bell, G., & Gonzalez, A. (2009). Evolutionary rescue can prevent extinction following environmental change. Ecology Letters, 12(9), 942–948. https://doi.org/10.1111/j.1461-0248.2009.01350.x

Bellard, C., Bertelsmeier, C., Leadley, P., Thuiller, W., & Courchamp, F. (2012). Impacts of climate change on the future of biodiversity. Ecology Letters, 15(4), 365–377. https://doi.org/10.1111/j.1461-0248.2011.01736.x

Benito Garzón, M., Robson, T. M., & Hampe, A. (2019). ΔTraitSDMs: Species distribution models that account for local adaptation and phenotypic plasticity. New Phytologist, 222(4), 1757–1765. https://doi.org/10.1111/nph.15716

Benjelloun, B., Boyer, F., Streeter, I., Zamani, W., Engelen, S., Alberti, A., Alberto, F. J., BenBati, M., Ibnelbachyr, M., Chentouf, M., Bechchari, A., Rezaei, H. R., Naderi, S., Stella, A., Chikhi, A., Clarke, L., Kijas, J., Flicek, P., Taberlet, P., & Pompanon, F. (2019). An evaluation of sequencing coverage and genotyping strategies to assess neutral and adaptive diversity. Molecular Ecology Resources, 19(6), 1497–1515. https://doi.org/10.1111/1755-0998.13070

Bittencourt-Silva, G. B., Lawson, L. P., Tolley, K. A., Portik, D. M., Barratt, C. D., Nagel, P., & Loader, S. P. (2017). Impact of species delimitation and sampling on niche models and phylogeographical inference: A case study of the East African reed frog Hyperolius substriatus Ahl, 1931. Molecular Phylogenetics and Evolution, 114, 261–270. https://doi.org/10.1016/j.ympev.2017.06.022

Brennan, A., Naidoo, R., Greenstreet, L., Mehrabi, Z., Ramankutty, N., & Kremen, C. (2022). Functional connectivity of the world’s protected areas. Science, 376(6597), 1101–1104. https://doi.org/10.1126/science.abl8974

Campbell, E. O., Brunet, B. M. T., Dupuis, J. R., & Sperling, F. A. H. (2018). Would an RRS by any other name sound as RAD? Methods in Ecology and Evolution, 9(9), 1920– 1927. https://doi.org/10.1111/2041-210X.13038

Capblancq, T., Fitzpatrick, M. C., Bay, R. A., Exposito-Alonso, M., & Keller, S. R. (2020). Genomic Prediction of (Mal)Adaptation Across Current and Future Climatic Landscapes. Annual Review of Ecology, Evolution, and Systematics, 51(1), 245–269. https://doi.org/10.1146/annurev-ecolsys-020720-042553

Capblancq, T., & Forester, B. R. (2021). Redundancy analysis: A Swiss Army Knife for landscape genomics. Methods in Ecology and Evolution, 12(12), 2298–2309. https://doi.org/10.1111/2041-210X.13722

Catchen, J. M., Hohenlohe, P. A., Bernatchez, L., Funk, W. C., Andrews, K. R., & Allendorf, F. W. (2017). Unbroken: RADseq remains a powerful tool for understanding the genetics of adaptation in natural populations. Molecular Ecology Resources, 17(3), 362–365. https://doi.org/10.1111/1755-0998.12669

Cerca, J., Maurstad, M. F., Rochette, N. C., Rivera-Colón, A. G., Rayamajhi, N., Catchen, J. M., & Struck, T. H. (2021). Removing the bad apples: A simple bioinformatic method to improve loci-recovery in de novo RADseq data for non-model organisms. Methods in Ecology and Evolution, 12(5), 805–817. https://doi.org/10.1111/2041-210X.13562

Chen, Y., Jiang, Z., Fan, P., Ericson, P. G. P., Song, G., Luo, X., Lei, F., & Qu, Y. (2022a). The combination of genomic offset and niche modelling provides insights into climate change-driven vulnerability. Nature Communications, 13(1), Article 1. https://doi.org/10.1038/s41467-022-32546-z

Chen, Z., Grossfurthner, L., Loxterman, J. L., Masingale, J., Richardson, B. A., Seaborn, T., Smith, B., Waits, L. P., & Narum, S. R. (2022b). Applying genomics in assisted migration under climate change: Framework, empirical applications, and case studies. Evolutionary Applications, 15(1), 3–21. https://doi.org/10.1111/eva.13335

Clark, L. V., Lipka, A. E., & Sacks, E. J. (2019). polyRAD: Genotype Calling with Uncertainty from Sequencing Data in Polyploids and Diploids. G3 Genes|Genomes|Genetics, 9(3), 663–673. https://doi.org/10.1534/g3.118.200913

Collart, F., Hedenäs, L., Broennimann, O., Guisan, A., & Vanderpoorten, A. (2021). Intraspecific differentiation: Implications for niche and distribution modelling. Journal of Biogeography, 48(2), 415–426. https://doi.org/10.1111/jbi.14009

Eaton, D. A. R., & Overcast, I. (2020). ipyrad: Interactive assembly and analysis of RADseq datasets. Bioinformatics, 36(8), 2592–2594. https://doi.org/10.1093/bioinformatics/btz966

Elith, J., & Leathwick, J. R. (2009). Species Distribution Models: Ecological Explanation and Prediction Across Space and Time. Annual Review of Ecology, Evolution, and Systematics, 40(1), 677–697. https://doi.org/10.1146/annurev.ecolsys.110308.120159

Exposito-Alonso, M., Booker, T. R., Czech, L., Gillespie, L., Hateley, S., Kyriazis, C. C., Lang, P. L. M., Leventhal, L., Nogues-Bravo, D., Pagowski, V., Ruffley, M., Spence, J. P., Toro Arana, S. E., Weiß, C. L., & Zess, E. (2022). Genetic diversity loss in the Anthropocene. Science, 377(6613), 1431–1435. https://doi.org/10.1126/science.abn5642

Exposito-Alonso, M., Vasseur, F., Ding, W., Wang, G., Burbano, H. A., & Weigel, D. (2018). Genomic basis and evolutionary potential for extreme drought adaptation in Arabidopsis thaliana. Nature Ecology & Evolution, 2(2), Article 2. https://doi.org/10.1038/s41559-017-0423-0

Fick, S. E., & Hijmans, R. J. (2017). WorldClim 2: New 1-km spatial resolution climate surfaces for global land areas. International Journal of Climatology, 37(12), 4302– 4315. https://doi.org/10.1002/joc.5086

Fitzpatrick, M. C., & Keller, S. R. (2015). Ecological genomics meets community-level modelling of biodiversity: Mapping the genomic landscape of current and future environmental adaptation. Ecology Letters, 18(1), 1–16. https://doi.org/10.1111/ele.12376

Foden, W. B., Young, B. E., Akçakaya, H. R., Garcia, R. A., Hoffmann, A. A., Stein, B. A., Thomas, C. D., Wheatley, C. J., Bickford, D., Carr, J. A., Hole, D. G., Martin, T. G., Pacifici, M., Pearce-Higgins, J. W., Platts, P. J., Visconti, P., Watson, J. E. M., & Huntley, B. (2019). Climate change vulnerability assessment of species. WIREs Climate Change, 10(1), e551. https://doi.org/10.1002/wcc.551

Forester, B. R., Lasky, J. R., Wagner, H. H., & Urban, D. L. (2018). Comparing methods for detecting multilocus adaptation with multivariate genotype–environment associations. Molecular Ecology, 27(9), 2215–2233. https://doi.org/10.1111/mec.14584

Forester, B. R., Beever, E. A., Darst, C., Szymanski, J., & Funk, W. C. (2022). Linking evolutionary potential to extinction risk: Applications and future directions. Frontiers in Ecology and the Environment, 20(9), 507–515. https://doi.org/10.1002/fee.2552

Forester, B. R., Day, C. C., Ruegg, K., & Landguth, E. L. (2023). Evolutionary potential mitigates extinction risk under climate change in the endangered southwestern willow flycatcher. Journal of Heredity, esac067. https://doi.org/10.1093/jhered/esac067

Fox, R. J., Donelson, J. M., Schunter, C., Ravasi, T., & Gaitán-Espitia, J. D. (2019). Beyond buying time: The role of plasticity in phenotypic adaptation to rapid environmental change. Philosophical Transactions of the Royal Society B: Biological Sciences, 374(1768), 20180174. https://doi.org/10.1098/rstb.2018.0174

Frankham, R. (2015). Genetic rescue of small inbred populations: Meta-analysis reveals large and consistent benefits of gene flow. Molecular Ecology, 24(11), 2610–2618. https://doi.org/10.1111/mec.13139

Frankham, R., Ballou, J. D., Ralls, K., Eldridge, M. D. B., Dudash, M. R., Fenster, C. B., Lacy, R. C., & Sunnucks, P. (2019). A practical guide for genetic management of fragmented animal and plant populations. Oxford University Press. https://doi.org/10.1093/oso/9780198783411.001.0001

Guisan, A., & Thuiller, W. (2005). Predicting species distribution: Offering more than simple habitat models. Ecology Letters, 8(9), 993–1009. https://doi.org/10.1111/j.1461-0248.2005.00792.x

Hällfors, M. H., Aikio, S., Fronzek, S., Hellmann, J. J., Ryttäri, T., & Heikkinen, R. K. (2016). Assessing the need and potential of assisted migration using species distribution models. Biological Conservation, 196, 60–68. https://doi.org/10.1016/j.biocon.2016.01.031

Hereford, J. (2009). A Quantitative Survey of Local Adaptation and Fitness Trade[Offs. The American Naturalist, 173(5), 579–588. https://doi.org/10.1086/597611

Hoban, S. (2014). An overview of the utility of population simulation software in molecular ecology. Molecular Ecology, 23(10), 2383–2401. https://doi.org/10.1111/mec.12741

Hoban, S., Archer, F. I., Bertola, L. D., Bragg, J. G., Breed, M. F., Bruford, M. W., Coleman, M. A., Ekblom, R., Funk, W. C., Grueber, C. E., Hand, B. K., Jaffé, R., Jensen, E., Johnson, J. S., Kershaw, F., Liggins, L., MacDonald, A. J., Mergeay, J., Miller, J. M., … Hunter, M. E. (2022). Global genetic diversity status and trends: Towards a suite of Essential Biodiversity Variables (EBVs) for genetic composition. Biological Reviews, 97(4), 1511–1538. https://doi.org/10.1111/brv.12852

Hoban, S., Campbell, C. D., da Silva, J. M., Ekblom, R., Funk, W. C., Garner, B. A., Godoy, J. A., Kershaw, F., MacDonald, A. J., Mergeay, J., Minter, M., O’Brien, D., Vinas, I. P., Pearson, S. K., Pérez-Espona, S., Potter, K. M., Russo, I.-R. M., Segelbacher, G., Vernesi, C., & Hunter, M. E. (2021). Genetic diversity is considered important but interpreted narrowly in country reports to the Convention on Biological Diversity: Current actions and indicators are insufficient. Biological Conservation, 261, 109233. https://doi.org/10.1016/j.biocon.2021.109233

Hoffmann, A. A., & Sgrò, C. M. (2011). Climate change and evolutionary adaptation. Nature, 470(7335), Article 7335. https://doi.org/10.1038/nature09670

Howard, C., Flather, C. H., & Stephens, P. A. (2020). A global assessment of the drivers of threatened terrestrial species richness. Nature Communications, 11(1), Article 1. https://doi.org/10.1038/s41467-020-14771-6

Ikeda, D. H., Max, T. L., Allan, G. J., Lau, M. K., Shuster, S. M., & Whitham, T. G. (2017). Genetically informed ecological niche models improve climate change predictions. Global Change Biology, 23(1), 164–176. https://doi.org/10.1111/gcb.13470

IPBES (2019). Global assessment report on biodiversity and ecosystem services of the Intergovernmental Science-Policy Platform on Biodiversity and Ecosystem Services. E. S. Brondizio, J. Settele, S. Díaz, and H. T. Ngo (editors). IPBES secretariat, Bonn, Germany

IPCC (2014). Climate Change 2014: Mitigation of Climate Change. Contribution of Working Group III to the Fifth Assessment Report of the Intergovernmental Panel on Climate Change [Edenhofer, O., R. Pichs-Madruga, Y. Sokona, E. Farahani, S. Kadner, K. Seyboth, A. Adler, I. Baum, S. Brunner, P. Eickemeier, B. Kriemann, J. Savolainen, S. Schlömer, C. von Stechow, T. Zwickel and J.C. Minx (eds.)]. Cambridge University Press, Cambridge, United Kingdom and New York, NY, USA

Johnston, A., Matechou, E., & Dennis, E. B. (2023). Outstanding challenges and future directions for biodiversity monitoring using citizen science data. Methods in Ecology and Evolution, 14(1), 103–116. https://doi.org/10.1111/2041-210X.13834

Karger, D. N., Conrad, O., Böhner, J., Kawohl, T., Kreft, H., Soria-Auza, R. W., Zimmermann, N. E., Linder, H. P., & Kessler, M. (2017). Climatologies at high resolution for the earth’s land surface areas. Scientific Data, 4(1), Article 1. https://doi.org/10.1038/sdata.2017.122

Korneliussen, T. S., Albrechtsen, A., & Nielsen, R. (2014). ANGSD: Analysis of Next Generation Sequencing Data. BMC Bioinformatics, 15(1), 356. https://doi.org/10.1186/s12859-014-0356-4

Kurtzer, G. M., Sochat, V., & Bauer, M. W. (2017). Singularity: Scientific containers for mobility of compute. PLOS ONE, 12(5), e0177459. https://doi.org/10.1371/journal.pone.0177459

Laikre, L., Schwartz, M. K., Waples, R. S., & Ryman, N. (2010). Compromising genetic diversity in the wild: Unmonitored large-scale release of plants and animals. Trends in Ecology & Evolution, 25(9), 520–529. https://doi.org/10.1016/j.tree.2010.06.013

Lancaster, L. T., Fuller, Z. L., Berger, D., Barbour, M. A., Jentoft, S., & Wellenreuther, M. (2022). Understanding climate change response in the age of genomics. Journal of Animal Ecology, 91(6), 1056–1063. https://doi.org/10.1111/1365-2656.13711

Layton, K. K. S., & Bradbury, I. R. (2022). Harnessing the power of multi-omics data for predicting climate change response. Journal of Animal Ecology, 91(6), 1064–1072. https://doi.org/10.1111/1365-2656.13619

Lehtomäki, J., & Moilanen, A. (2013). Methods and workflow for spatial conservation prioritization using Zonation. Environmental Modelling & Software, 47, 128–137. https://doi.org/10.1016/j.envsoft.2013.05.001

Lowry, D. B., Hoban, S., Kelley, J. L., Lotterhos, K. E., Reed, L. K., Antolin, M. F., & Storfer, A. (2017). Breaking RAD: An evaluation of the utility of restriction site-associated DNA sequencing for genome scans of adaptation. Molecular Ecology Resources, 17(2), 142–152. https://doi.org/10.1111/1755-0998.12635

Maxwell, S. L., Fuller, R. A., Brooks, T. M., & Watson, J. E. M. (2016). Biodiversity: The ravages of guns, nets and bulldozers. Nature, 536(7615), Article 7615. https://doi.org/10.1038/536143a

McGuire, J. L., Lawler, J. J., McRae, B. H., Nuñez, T. A., & Theobald, D. M. (2016). Achieving climate connectivity in a fragmented landscape. Proceedings of the National Academy of Sciences, 113(26), 7195–7200. https://doi.org/10.1073/pnas.1602817113

McKenna, A., Hanna, M., Banks, E., Sivachenko, A., Cibulskis, K., Kernytsky, A., Garimella, K., Altshuler, D., Gabriel, S., Daly, M., & DePristo, M. A. (2010). The Genome Analysis Toolkit: A MapReduce framework for analyzing next-generation DNA sequencing data. Genome Research, 20(9), 1297–1303. https://doi.org/10.1101/gr.107524.110

Merila, J., & Hendry, A.P. (2014). Climate change, adaptation, and phenotypic plasticity: the problem and the evidence. Evolutionary Applications, 7(1), 1–14. https://doi.org/10.1111/eva.12137

Merow, C., Smith, M. J., & Silander Jr, J. A. (2013). A practical guide to MaxEnt for modeling species’ distributions: What it does, and why inputs and settings matter. Ecography, 36(10), 1058–1069. https://doi.org/10.1111/j.1600-0587.2013.07872.x

Moilanen, A., Franco, A. M. A., Early, R. I., Fox, R., Wintle, B., & Thomas, C. D. (2005). Prioritizing multiple-use landscapes for conservation: Methods for large multi-species planning problems. Proceedings of the Royal Society B: Biological Sciences, 272(1575), 1885–1891. https://doi.org/10.1098/rspb.2005.3164

Nadeau, C. P., & Urban, M. C. (2019). Eco-evolution on the edge during climate change. Ecography, 42(7), 1280–1297. https://doi.org/10.1111/ecog.04404

Oomen, R. A., & Hutchings, J. A. (2022). Genomic reaction norms inform predictions of plastic and adaptive responses to climate change. Journal of Animal Ecology, 91(6), 1073–1087. https://doi.org/10.1111/1365-2656.13707

Pacifici, M., Foden, W. B., Visconti, P., Watson, J. E. M., Butchart, S. H. M., Kovacs, K. M., Scheffers, B. R., Hole, D. G., Martin, T. G., Akçakaya, H. R., Corlett, R. T., Huntley, B., Bickford, D., Carr, J. A., Hoffmann, A. A., Midgley, G. F., Pearce-Kelly, P., Pearson, R. G., Williams, S. E., … Rondinini, C. (2015). Assessing species vulnerability to climate change. Nature Climate Change, 5(3), Article 3. https://doi.org/10.1038/nclimate2448

Paris, J. R., Stevens, J. R., & Catchen, J. M. (2017). Lost in parameter space: A road map for stacks. Methods in Ecology and Evolution, 8(10), 1360–1373. https://doi.org/10.1111/2041-210X.12775

Parks, S. A., Holsinger, L. M., Littlefield, C. E., Dobrowski, S. Z., Zeller, K. A., Abatzoglou, J. T., Besancon, C., Nordgren, B. L., & Lawler, J. J. (2022). Efficacy of the global protected area network is threatened by disappearing climates and potential transboundary range shifts. Environmental Research Letters, 17(5), 054016. https://doi.org/10.1088/1748-9326/ac6436

Parmesan, C. (2006). Ecological and Evolutionary Responses to Recent Climate Change. Annual Review of Ecology, Evolution, and Systematics, 37(1), 637–669. https://doi.org/10.1146/annurev.ecolsys.37.091305.110100

Patton, A. H., Margres, M. J., Stahlke, A. R., Hendricks, S., Lewallen, K., Hamede, R. K., Ruiz-Aravena, M., Ryder, O., McCallum, H. I., Jones, M. E., Hohenlohe, P. A., & Storfer, A. (2019). Contemporary Demographic Reconstruction Methods Are Robust to Genome Assembly Quality: A Case Study in Tasmanian Devils. Molecular Biology and Evolution, 36(12), 2906–2921. https://doi.org/10.1093/molbev/msz191

Pauls, S. U., Nowak, C., Bálint, M., & Pfenninger, M. (2013). The impact of global climate change on genetic diversity within populations and species. Molecular Ecology, 22(4), 925–946. https://doi.org/10.1111/mec.12152

Pecl, G. T., Araújo, M. B., Bell, J. D., Blanchard, J., Bonebrake, T. C., Chen, I.-C., Clark, T. D., Colwell, R. K., Danielsen, F., Evengård, B., Falconi, L., Ferrier, S., Frusher, S., Garcia, R. A., Griffis, R. B., Hobday, A. J., Janion-Scheepers, C., Jarzyna, M. A., Jennings, S., … Williams, S. E. (2017). Biodiversity redistribution under climate change: Impacts on ecosystems and human well-being. Science, 355(6332), eaai9214. https://doi.org/10.1126/science.aai9214

Peterman, W. E. (2018). ResistanceGA: An R package for the optimization of resistance surfaces using genetic algorithms. Methods in Ecology and Evolution, 9(6), 1638–1647. https://doi.org/10.1111/2041-210X.12984

Pimsler, M. L., Oyen, K. J., Herndon, J. D., Jackson, J. M., Strange, J. P., Dillon, M. E., & Lozier, J. D. (2020). Biogeographic parallels in thermal tolerance and gene expression variation under temperature stress in a widespread bumble bee. Scientific Reports, 10(1), Article 1. https://doi.org/10.1038/s41598-020-73391-8

Pinsky, M. L., Comte, L., & Sax, D. F. (2022). Unifying climate change biology across realms and taxa. Trends in Ecology & Evolution, 37(8), 672–682. https://doi.org/10.1016/j.tree.2022.04.011

Pinsky, M. L., Eikeset, A. M., McCauley, D. J., Payne, J. L., & Sunday, J. M. (2019). Greater vulnerability to warming of marine versus terrestrial ectotherms. Nature, 569(7754), Article 7754. https://doi.org/10.1038/s41586-019-1132-4

Pfenninger, M., & Bálint, M. (2022). On the use of population genomic time series for environmental monitoring. American Journal of Botany, 109(4), 497–499. https://doi.org/10.1002/ajb2.1836

Pfenninger, M., Foucault, Q., Waldvogel, A.-M., & Feldmeyer, B. (2023). Selective effects of a short transient environmental fluctuation on a natural population. Molecular Ecology, 32(2), 335–349. https://doi.org/10.1111/mec.16748

Purcell, S., Neale, B., Todd-Brown, K., Thomas, L., Ferreira, M. A. R., Bender, D., Maller, J., Sklar, P., Bakker, P. I. W. de, Daly, M. J., & Sham, P. C. (2007). PLINK: A Tool Set for Whole-Genome Association and Population-Based Linkage Analyses. The American Journal of Human Genetics, 81(3), 559–575. https://doi.org/10.1086/519795

Puritz, J. B., Hollenbeck, C. M., & Gold, J. R. (2014). dDocent: A RADseq, variant-calling pipeline designed for population genomics of non-model organisms. PeerJ, 2, e431. https://doi.org/10.7717/peerj.431

Quintero, I., & Wiens, J. J. (2013). Rates of projected climate change dramatically exceed past rates of climatic niche evolution among vertebrate species. Ecology Letters, 16(8), 1095–1103. https://doi.org/10.1111/ele.12144

Radchuk, V., Reed, T., Teplitsky, C., van de Pol, M., Charmantier, A., Hassall, C., Adamík, P., Adriaensen, F., Ahola, M. P., Arcese, P., Miguel Avilés, J., Balbontin, J., Berg, K. S., Borras, A., Burthe, S., Clobert, J., Dehnhard, N., de Lope, F., Dhondt, A. A., … Kramer-Schadt, S. (2019). Adaptive responses of animals to climate change are most likely insufficient. Nature Communications, 10(1), Article 1. https://doi.org/10.1038/s41467-019-10924-4

Razgour, O., Forester, B., Taggart, J. B., Bekaert, M., Juste, J., Ibáñez, C., Puechmaille, S. J., Novella-Fernandez, R., Alberdi, A., & Manel, S. (2019). Considering adaptive genetic variation in climate change vulnerability assessment reduces species range loss projections. Proceedings of the National Academy of Sciences, 116(21), 10418– 10423. https://doi.org/10.1073/pnas.1820663116

Razgour, O., Taggart, J. B., Manel, S., Juste, J., Ibáñez, C., Rebelo, H., Alberdi, A., Jones, G., & Park, K. (2018). An integrated framework to identify wildlife populations under threat from climate change. Molecular Ecology Resources, 18(1), 18–31. https://doi.org/10.1111/1755-0998.12694

Rellstab, C., Dauphin, B., & Exposito-Alonso, M. (2021). Prospects and limitations of genomic offset in conservation management. Evolutionary Applications, 14(5), 1202– 1212. https://doi.org/10.1111/eva.13205

Ruegg, K., Bay, R. A., Anderson, E. C., Saracco, J. F., Harrigan, R. J., Whitfield, M., Paxton, E. H., & Smith, T. B. (2018). Ecological genomics predicts climate vulnerability in an endangered southwestern songbird. Ecology Letters, 21(7), 1085–1096. https://doi.org/10.1111/ele.12977

Schielzeth, H., & Wolf, J. B. W. (2021). Community genomics: A community-wide perspective on within-species genetic diversity. American Journal of Botany, 108(11), 2108–2111. https://doi.org/10.1002/ajb2.1796

Schipper, A. M., Hilbers, J. P., Meijer, J. R., Antão, L. H., Benítez-López, A., de Jonge, M. M. J., Leemans, L. H., Scheper, E., Alkemade, R., Doelman, J. C., Mylius, S., Stehfest, E., van Vuuren, D. P., van Zeist, W.-J., & Huijbregts, M. A. J. (2020). Projecting terrestrial biodiversity intactness with GLOBIO 4. Global Change Biology, 26(2), 760–771. https://doi.org/10.1111/gcb.14848

Segelbacher, G., Bosse, M., Burger, P., Galbusera, P., Godoy, J. A., Helsen, P., Hvilsom, C., Iacolina, L., Kahric, A., Manfrin, C., Nonic, M., Thizy, D., Tsvetkov, I., Veličković, N., Vilà, C., Wisely, S. M., & Buzan, E. (2022). New developments in the field of genomic technologies and their relevance to conservation management. Conservation Genetics, 23(2), 217–242. https://doi.org/10.1007/s10592-021-01415-5

Smith, T. B., Fuller, T. L., Zhen, Y., Zaunbrecher, V., Thomassen, H. A., Njabo, K., Anthony, N. M., Gonder, M. K., Buermann, W., Larison, B., Ruegg, K., & Harrigan, R. J. (2021). Genomic vulnerability and socio-economic threats under climate change in an African rainforest bird. Evolutionary Applications, 14(5), 1239–1247. https://doi.org/10.1111/eva.13193

Stange, M., Barrett, R. D. H., & Hendry, A. P. (2021). The importance of genomic variation for biodiversity, ecosystems and people. Nature Reviews Genetics, 22(2), Article 2. https://doi.org/10.1038/s41576-020-00288-7

Taus, T., Futschik, A., & Schlötterer, C. (2017). Quantifying Selection with Pool-Seq Time Series Data. Molecular Biology and Evolution, 34(11), 3023–3034. https://doi.org/10.1093/molbev/msx225

Teixeira, J. C., & Huber, C. D. (2021). The inflated significance of neutral genetic diversity in conservation genetics. Proceedings of the National Academy of Sciences, 118(10), e2015096118. https://doi.org/10.1073/pnas.2015096118

Thuiller, W., Lafourcade, B., Engler, R., & Araújo, M. B. (2009). BIOMOD – a platform for ensemble forecasting of species distributions. Ecography, 32(3), 369–373. https://doi.org/10.1111/j.1600-0587.2008.05742.x

Tilman, D., Clark, M., Williams, D. R., Kimmel, K., Polasky, S., & Packer, C. (2017). Future threats to biodiversity and pathways to their prevention. Nature, 546(7656), Article 7656. https://doi.org/10.1038/nature22900

Urban, M. C. (2015). Accelerating extinction risk from climate change. Science, 348(6234), 571–573. https://doi.org/10.1126/science.aaa4984

Waldvogel, A.-M., Feldmeyer, B., Rolshausen, G., Exposito-Alonso, M., Rellstab, C., Kofler, R., Mock, T., Schmid, K., Schmitt, I., Bataillon, T., Savolainen, O., Bergland, A., Flatt, T., Guillaume, F., & Pfenninger, M. (2020). Evolutionary genomics can improve prediction of species’ responses to climate change. Evolution Letters, 4(1), 4–18. https://doi.org/10.1002/evl3.154

Waldvogel, A.-M., Schreiber, D., Pfenninger, M., & Feldmeyer, B. (2020). Climate Change Genomics Calls for Standardized Data Reporting. Frontiers in Ecology and Evolution, 8. https://www.frontiersin.org/articles/10.3389/fevo.2020.00242

Wollenberg Valero, K. C., Garcia-Porta, J., Irisarri, I., Feugere, L., Bates, A., Kirchhof, S., Jovanović Glavaš, O., Pafilis, P., Samuel, S. F., Müller, J., Vences, M., Turner, A. P., Beltran-Alvarez, P., & Storey, K. B. (2022). Functional genomics of abiotic environmental adaptation in lacertid lizards and other vertebrates. Journal of Animal Ecology, 91(6), 1163–1179. https://doi.org/10.1111/1365-2656.13617

Zizka, A., Silvestro, D., Andermann, T., Azevedo, J., Duarte Ritter, C., Edler, D., Farooq, H., Herdean, A., Ariza, M., Scharn, R., Svantesson, S., Wengström, N., Zizka, V., & Antonelli, A. (2019). CoordinateCleaner: Standardized cleaning of occurrence records from biological collection databases. Methods in Ecology and Evolution, 10(5), 744–751. https://doi.org/10.1111/2041-210X.13152

Zurell, D., Franklin, J., König, C., Bouchet, P. J., Dormann, C. F., Elith, J., Fandos, G., Feng, X., Guillera-Arroita, G., Guisan, A., Lahoz-Monfort, J. J., Leitão, P. J., Park, D. S., Peterson, A. T., Rapacciuolo, G., Schmatz, D. R., Schröder, B., Serra-Diaz, J. M., Thuiller, W., … Merow, C. (2020). A standard protocol for reporting species distribution models. Ecography, 43(9), 1261–1277. https://doi.org/10.1111/ecog.04960

