## Supporting Information 1 for "Life on the edge: a new toolbox for population-level climate change vulnerability assessments"

**Supplementary Text S1**. Details of LotE input data, functions and how output metrics are quantified

*Data preparation*

**prepare_spatial_data()** prepares the spatial data prior to building SDMs. Briefly, the script will download existing GBIF data, clean it (Zizka et al. 2019) and combine with the existing georeferenced genomic data, as well as cropping present and future climate data to a buffer around all presences. The *params* file may be populated with user specific requirements for each dataset, including a user-defined geographic extent, size of buffer to crop data (Table S1). **prepare_environmental_data()** prepares the environmental data that you have already downloaded (e.g. Worldclim2 or CHELSA), crops it to the extent for your study species and places it in its own directory within the species folder in *-data-*. Additionally, environmental predictor data will be extracted for each sampled individual based on the georeferenced coordinates of the genomic data for use with later GEA analyses.

**spatially_rarefy_presences_and_create_background_data()** will reduce spatial autocorrelation in the input presence data based on a distance defined in the params file Params file variables of relevance are spatial rarefaction distance buffer, and number of background points to select.

*Exposure*

**sdms_biomod2()** will use the spatially rarefied presence data to build species distribution models (SDMs) for each species, for present conditions, but also for forecasted future conditions. Here, the framework follows SDM best practices (see Araujo et al. 2019), using reduced spatial autocorrelation in presence data, as well as accounting for multi-collinearity in predictor variables using Variance Inflation Factors (VIF) if defined in the *params* file. Alternatively, users may specify a subset of predictors that are ecologically relevant for the species in question. The biomod2 R package (Thuiller et al. 2009) is used to evaluate models built using available modelling algorithms which is also subsettable via the params file), and retaining only ‘good’ models (i.e. TSS>0.5, modifiable in the params file) for the final ensemble species distribution model prediction (see Araújo & New, 2007). The model will then be projected onto the future climatic predictions to forecast the species distribution in the future. Variable importances will also be tracked across all retained models so that the user has a sense of which predictor variables have the most influence on the species distribution. Several SDM parameters are modifiable in the *params* file, including if VIF should be used, or a subset of variables should be selected, which SDM algorithms to use, number of replicates per algorithm, data split percentage between model training and testing, minimum ROC/TSS values to retain model replicates for ensemble modelling (Table S1). **exposure()** compares how selected climate variables and SDM suitability change between the present and predicted future conditions. Default behaviour is to use deciles of the minimum and maximum environmental extremes to assign dissimilarity scores (ranging between 1, minimal change and 10, the most change) for each predictor and the SDMs, though this can be modified in the params file (see Table S1 for details). A combined metric for ‘**Exposure**’ is then quantified, measuring the magnitude of change each population/locality is predicted to experience between the two time periods. In the *params* file the user can assign the relevant environmental predictors included in the exposure calculations (i.e. the same variables that will be used for GEAs). Exposure ranges from 0-10 and is calculated depending on the exposure rule variable in the *params* file, which can be used to modify how exposure is calculated by weighting the SDM and environmental predictor contributions, see Table S1. **impute_missing_data()** uses the *.ped* and *.map* files. Firstly, some file conversions will occur (into *.geno* and *.lfmm* formats), and a simple population structure analysis will be performed using sNMF (Frichot et al. 2014) to calculate the most likely number of populations ancestry co-efficients. The script will automatically select the number of populations represented by the data (k) with the lowest cross-entropy. Secondly, a PCA analysis (Oksanen et al. 2012) on the same data will be used to visualize population structure in ordination space. Before going further with the Genotype-Environment Association analyses, the output plots need to be checked, before as population structure can be a confounding factor in those analyses and needs to be accounted for to gain reliable results. A final important step before the next functions is to impute missing data, which will be done in two ways, firstly using LEA (Frichot & François, 2015), and secondly based on the mean frequencies of genotypes per population cluster (see Razgour et al. 2018). The *params* file allows the min and max iterations of k to test to be set, along with the number of sNMF replicates to run and the ploidy of the data.

*Sensitivity*

**gea_lfmm()** runs the first of the two Genotype-Environment Association analyses in this toolbox using LFMM (Caye et al. 2019), which is a univariate method. Firstly, the script will read in the environmental data as well as the imputed genomic data. The first iteration of the GEA will be run using LFMM – the user needs to define the number of populations (k) in the *params* file based on the population structure results generated by impute_missing_data(). The user should inspect the p-value distributions in the output plots generated for each predictor variable to ensure they are satisfied with the False Discovery Rate (FDR). If adjustments are needed (which is likely), the Genomic Inflation Factor (GIF) should be set in the *params* file (see Table S1), and the function can be re-run. The GIF represents inflation of test scores due to confounding factors in the data that have not been accounted for in the model, and so p-values, when adjusted, become q-values.. The function will automatically use the associated q-values set to select putative candidate SNPs at FDR thresholds of 0.01,

0.05 and 0.1, and output summary .csv files of the adaptive SNPs. **gea_rda()** is the second of the GEA analyses used in this toolbox, using Redundancy Analysis (RDA) from the vegan R package (Oksanen et al. 2012), which is a multivariate method. The script here follows a similar process to the LFMM analyses, reading in the environmental and genomic data before performing the initial RDA. A plot will be made to summarise the RDA and where each genetic cluster falls in the ordination space in relation to the environmental predictors, as well as a screeplot with the eigenvalues from each RDA axis. For post-processing of the RDA results based on the test scores, two approaches will be used. Firstly, the distribution of the SNPs will be categorised and outliers with a defined standard deviation from the mean of the RDA loadings will be used to threshold SNPs as candidates (Forester et al. 2018, Razgour et al. 2019). Secondly, the same approach as used for the LFMM script above can be used, by modifying the GIF and inspecting p-value distributions until satisfactory distributions per predictor and FDRs are obtained. Parameterisable options in the *params* file include the number of RDA axes to retain, the scaling factor for the GIF, or the minimum standard deviation from the mean for the SNPs to be classed as candidates.

**gea_rda_individual_categorisation()** categorises each individual sample based on their relative position in the constrained RDA ordination space relative to the environmental predictors (see Razgour et al. 2019). The script automatically reads in putative candidate SNPs that have been identified by either/both LFMM and RDA (definable in params file) and then performs a new RDA based on only these candidate SNPs and the environmental data. The built-in functions automatically select individuals that are falling within the range of certain conditions following Razgour et al. (2019). For example, in a two predictor model (e.g. rainfall and temperature), certain individuals may be classed as hot-dry or cold-wet adapted. Individuals that fall in between the ordination categorisations of either of these categories may be categorised as ‘intermediate’. The distribution of these categorised individuals is plotted in the ordination space, and each individual, which category it belongs to, and its geographic location is saved in a .csv file. Categories (e.g. 'hot-dry'/'cold-wet') may be defined in the *params* file, in addition to selecting which candidates are retained across the RDA and LFMM analyses (see vignette and Table S1). **adaptive_diversity()** quantifies the adaptive diversity across all populations by plotting the proportions of individuals that are putatively adapted to each category, in that their genotypes after individual categorization fall into the categories defined in the params file (e.g. hot-dry/cold-wet/intermediate). It uses the outputs generated by the individual categorisation (above), and plots a summary map of all samples (individuals and populations) and which conditions they are putatively adapted to. **neutral_diversity()** calculates neutral (i.e. non-adaptive) genetic diversity by masking out the putatively adaptive SNPs. PLINK will be used to generate the output files and the script here will automatically count the populations, number of individuals and neutral heterozygosity (excluding monomorphic SNPs across all populations) to calculate '**Neutral sensitivity**' (ranging from 1-10). Similar to quantifying exposure, default behaviour is to use deciles of the minimum and maximum extremes (of the data, in this case, observed heterozygosity) to assign neutral sensitivity metrics for each population, though this can be modified in the params file (see Table S1 for details). **sensitivity()** integrates the calculated neutral diversity and exposure with the proportions of adaptively categorised individuals across each population. Based on these an '**Adaptive sensitivity**' metric will be generated per population (i.e. unique geographic location) based on the % predicted change in each predictor (between current and future time periods) and the proportion of individuals locally adapted to that predictor following the below equation:

$$\text{Adaptive sensitivity} = \begin{array}{l}\text{Change in predictor 1 *}\\ \text{proportion of individuals}\\ \text{adapted to predictor 1}\end{array} + \begin{array}{l}\text{Change in predictor 2 *}\\ \text{proportion of individuals}\\ \text{adapted to predictor 2}\end{array}$$

The adaptive sensitivity metric will range between 1-10; being lower if a population has many individuals adapted to a condition that is forecast to change minimally, and higher if the conditions are forecast to change substantially. As an additional feature, local adaptation results may be used to predict the range shifts of individuals and populations with locally adapted genotypes, following the framework of Razgour et al. (2019) so that predicted range shifts for particular adaptive categories can be visualised. In this case, **adaptive_sdms()** builds new SDMs that parse individuals in the genomic dataset into their respective categories. It will re-perform the SDMs but instead of using the whole species presence data as input it will use each categorised *.csv* file generated by the gea_rda_individual_categorisation() function. Analytical details for the SDMs will follow the parameters defined in the *params* file.

*Range shift potential*

**create_circuitscape_inputs()** will set up the necessary files and structure for the range shift potential analyses. Circuitscape (Anantharaman et al. 2020) will be used to model predicted connectivity between populations in geographic space based on electrical circuit theory, based on a cumulative input resistance surface generated by the user (high values = more landscape resistance, low values = less resistance). This resistance surface can be generated by LotE by supplying a list of the selected variables to consider (taken from the spatial data folder) and their relative weights to contribute to the cumulative surface (both in the *params* file, see Table S1), or can be prepared outside of the toolbox (e.g. using software such as ResistanceGA, Peterman 2018) and added manually (see Fig. 2 for details). Because the analyses will model pairwise connectivity, memory and time requirements can become substantial with large datasets (i.e many populations/localities). This script will write the required files (*.ini, .jl*) for Circuitscape to run in Julia, along with the necessary files (*.sh*) to submit these jobs via HPC. The script writes relevant files to the Circuitscape directory for analysis as well as a summary of the resistance layers, and generates the specific required file for the spatial points input (derived from the input spatial genomic data), which is used to specify the 'nodes' to model connectivity between populations. In the *params* file, an option to transform all 0's to values of 0.001 is provided (recommended if running range shift potential analyses on SDM outputs for example) so that Circuitscape does not interpret unsuitable areas as completely impermeable barriers. The LotE Toolbox will run Circuitscape using Julia and the specific files that have been prepared. **range_shift_potential()** reads in the output data from the Circuitscape analyses, normalises the values so that they are between 0 (low) and 1 (high) based on the minimum and maximum values from the data, and plots an output map of modelled landscape  connectivity. It extracts the mean (normalised) connectivity of reach population to all other populations within a maximum dispersal distance (defined for each species in the params file, i.e. this avoids unrealistically distant populations being considered when calculating mean connectivity for each population), and then uses this to calculate '**Range shift potential**' (1-10), high reduction in range shift potential from present-future = 10, minimal reduction, stable or increase in range shift potential = 1.

*Population vulnerability*

**population_vulnerability()** integrates the exposure, adaptive and neutral sensitivity and range shift potential results to create a summary **'Population vulnerability**' metric per population ranging from low (1) to high (10).

The default behaviour is to to quantify population vulnerability for each population (i.e. unique geographic location) by taking the mean of the exposure, neutral sensitivity, adaptive sensitivity and range shift potential metrics as below, though this can be modified in the params file with the vulnerability_rule variable to weight the contributions of particular metrics if desired (see Table S1 for details):

$$\text{Population vulnerability} = \frac{(\text{Exposure} + \text{neutral sensitivity} + \text{adaptive sensitivity} + \text{range shift potential})}{4}$$

The function will create a *.csv* output file of the observed data (i.e. where genomic samples are located, and use an inverse distance weighted interpolation to predict Exposure, neutral sensitivity, adaptive sensitivity and Range shift potential results across geographical space, with the underlying assumption that locations closer together share similar properties than those further apart. The function produces summary maps of non-interpolated (observed) and interpolated (predicted) data for each of the metrics (separately and also a composite 4-panel map, see ). The *params* file allows the user to decide how population vulnerability is calculated by weighting the exposure, adaptive and neutral sensitivity and range shift potential metrics. **summary_pdfs()** uses all outputs generated and information in the log file to paste results together into a final summary PDF sheet using the grobblR package (Floyd, 2020). This, along with the extensive information recorded in the log file when each script has run provides a detailed summary of the steps of every analysis for the LotE Toolbox for each species. Results can be identified and probed by rerunning the individual functions with modified parameter settings. We recommend thorough reporting and transparency in all publications that use this toolbox, and urge users to provide log files, params files and PDF summaries as well as original input files.

*Supplementary Figures*

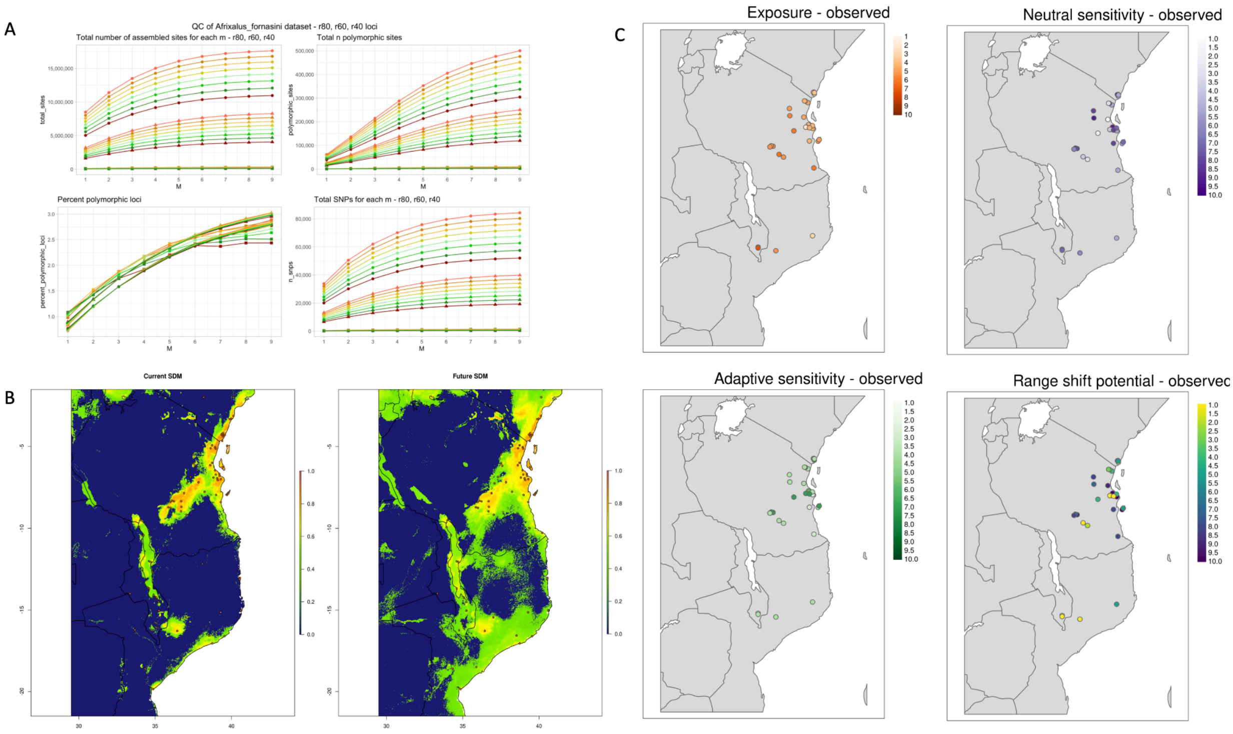


**Fig. S1.** *Afrixalus fornasini results.* A) Optimisation of Stacks parameters to assess total number of assembled SNPs, number of polymorphic SNPs across 40, 60 and 80% complete data matrices. A total of 72 different parameter combinations were tested (see main text for details). B) Current and future ensemble SDMs. C) Summary of observed LotE outputs per population for Exposure, neutral and adaptive Sensitivity, and Range shift potential.

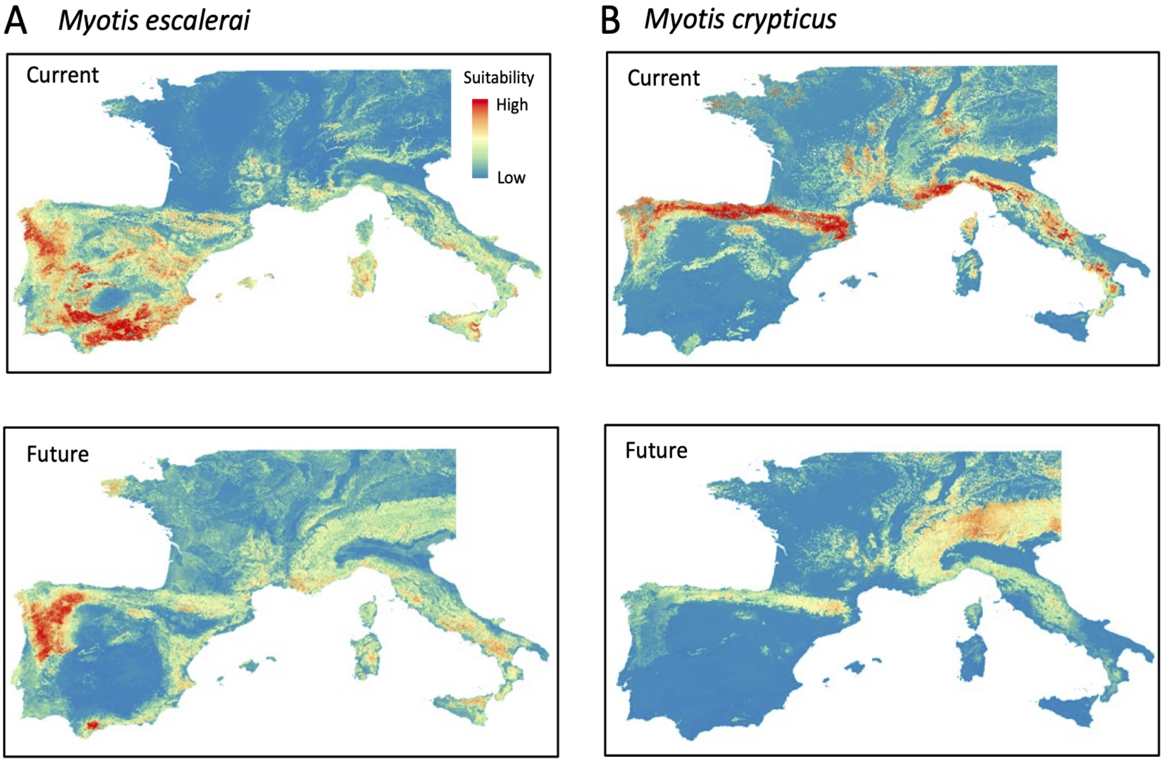


**Fig. S2.** A) *M. escalerai* present and future SDM ensembles, B) *M. crypticus* present and future SDM ensembles.

*Supplementary Tables*

**Table S1.** Modifiable toolbox parameters with description.

| Parameter name | Description |
|---|---|
| species_binomial | species name |
| user_specified_presences | a file containing presence records of the species (circumvents the need for obtaining data from GBIF). Useful if you have a species with limited publicly available data. The .csv formatted file should contain 2 columns named 'LONG', and 'LAT' |
| gbif_limit | maximum number of records to download from GBIF database, useful particularly for common species that may have thousands to millions of records |
| gbif_date_limit_lower | earliest date to retain GBIF records from (4 digits, e.g. 1990), to be used with gbif_date_limit_upper |
| gbif_date_limit_upper | latest date to retain GBIF records from (4 digits, e.g. 2023), to be used with gbif_date_limit_lower |
| climate_res | resolution of predictor data when spatially rarefying background points to speed up the process in case you have high resolution data |
| geographic_extent | define geographic area if desired (xmin,xmax,ymin,ymax) |
| current_climate_data_path | directory with current climate data used to clip extent for each study species |
| future_climate_data_path | directory with future climate data used to clip extent for each study species |
| sp_rare_dist_km | spatial distance (km) to rarefy samples |
| sdm_reps_per_algorithm | number of replicates per SDM algorithm to use |
| bg_buff_dist_degrees | number of decimal degrees buffer around presence points to use when generating SDM data for background points |
| n_background_points | number of background points to randomly sample in background buffer |
| perform_vif | automatically remove SDM predictor variables that have a VIF of >10 (requires at least 3 variables to be used) |
| subset_predictors | accepts a subset list of predictors separated by quotes and commas to use for SDMs if required (e.g. 'bioclim_1, bioclim_2'). Note that perform_vif should be set to no if this option is used |
| biomod_algorithms | accepts a subset of biomod2 SDM algorithms separated by quotes and commas to run if required (e.g. 'MAXENT', 'RF') |
| maxent_path | path to maxent.jar file (used for biomod2) |
| data_split_percentage | percentage of the data to be taken for training vs. testing in SDMs |
| AUC_min | minimum AUC score required to take model further into ensemble modelling |
| TSS_min | minimum TSS score required to take model further into ensemble modelling |
| env_predictor_1 | first environmental predictor to use for GEAs |
| env_predictor_2 | second environmental predictor to use for GEAs |
| exposure_quantification | used to decide how to assign exposure values - 'interval' will divide the extracted predictor values for each population (SDM, env_1, env_2) into 10 equal measures and assign an exposure score from low (1) - high (10) depending on the magnitude of change. 'quantile' will use the quantiles of the data (based on fitting the data into a normal distribution), 'defined' will use a predefined scale of 10 values specified by the user for |

| | |
|---|---|
| | each of the exposure_sdm_threshold, exposure_env_1_threshold, exposure_env_2_threshold variables. If studying a single species use either 'interval' or 'quantile', if multiple species we recommend using the 'defined' option so that results across species datasets are comparable |
| exposure_sdm_thresholds | string of 10 variables separated by comma for defining the exposure of SDM change |
| exposure_env_1_thresholds | string of 10 variables separated by comma for defining the exposure of env_1 |
| exposure_env_2_thresholds | string of 10 variables separated by comma for defining the exposure of env_2 |
| k_min | minimum number of K (=populations) to iterate population structure analyses |
| k_max | maximum number of K (=populations) to iterate population structure analyses |
| snmf_reps | number of sNMF replications to iterate each value of k |
| ploidy | ploidy of the species |
| plink_executable | path to plink binaries |
| neutral_sensitivity_quantification | used to decide how to assign neutral sensitivity values - 'interval' will divide the extracted neutral heterozygosity values for each population into 10 equal measures and assign a score from low (1) - high (10) depending on the values. 'quantile' will use the quantiles of the data (based on fitting the data into a normal distribution), 'defined' will use a predefined scale of 10 values specified by the user for each of the neutral_sensitivity_threshold values. If studying a single species use either 'interval' or 'quantile', if multiple species we recommend using the 'defined' option so that results across species datasets are comparable |
| neutural_sensitivity_thresholds | string of 10 variables separated by comma for defining the neutral sensitivity of populations |
| k | number of k to use after evaluating population structure analyses |
| scale_gif_lfmm | scaling value to modify Genomic Inflation Factor for LFMM analyses |
| rda_axes | number of axes to retain in RDA analyses |
| rda_SD | standard deviation to use as a cutoff to determine outliers for RDA analyses |
| scale_gif_rda | scaling value to modify Genomic Inflation Factor for RDA analyses |
| which_loci | which loci to retain for subsequent analyses: # 1: present in either RDA (SD<3) or LFMM (FDR<0.05), # 2: present in either RDA (FDR<0.05) or LFMM (FDR<0.05), # 3: only LFMM loci (FDR<0.05), # 4: only RDA loci (SD<3), # 5: only RDA loci (FDR<0.05), # 6: present in both RDA (SD<3) and LFMM (FDR<0.05), # 7: present in both RDA (FDR<0.05) and LFMM (FDR<0.05) - not recommended, often 0 loci |
| adaptive_sensitivity_quantification | used to decide how to assign adaptive sensitivity values - 'interval' will divide the extracted adaptive diversity values for each population into 10 equal measures and assign a score from low (1) - high (10) depending on the values. 'quantile' will use the quantiles of the data (based on fitting the data into a normal distribution), 'defined' will use a predefined scale of 10 values specified by the user for each of the adaptive_sensitivity_threshold values. If studying a single species use either 'interval' or 'quantile', if multiple species we recommend using the 'defined' option so that results across species datasets are comparable |
| adaptive_sensitivity_thresholds | string of 10 variables separated by comma for defining the adaptive sensitivity of populations |
| category_1 | name of adaptive category 1 to classify samples based on position in RDA ordination space |
| category_2 | name of adaptive category 2 to classify samples based on position in RDA ordination space |
| circuitscape_transform_zeros | Transform 0 values to 0.01 in input layers so that they are not complete barriers |
| circuitscape_layers | list of the input layers used to parameterise a cumulative landscape resistance surface |

| | |
|---|---|
| circuitscape_weights | relative weights of each of the circuitscape layers - the numbers supplied should be comma separated, adding to a total of 1, and match the number of circuitscape layers |
| circuitscape_max_dispersal_distance_km | maximum dispersal distance (in km) for the species used for calculating mean connectivity in range shift potential analyses |
| exposure_rule | rule for calculating exposure: # 1: Exposure = mean score of sdm and environmental predictors, # 2: Exposure = 25% SDM, 75% environmental predictors, # 3: Exposure = 50% SDM, 50% environmental predictors, # 4: Exposure = 75% SDM, 25% environmental predictors |
| population_vulnerability_quantification | used to decide how to assign population vulnerability values - 'interval' will divide the extracted population vulnerability values for each population into 10 equal measures and assign a score from low (1) - high (10) depending on the values. 'quantile' will use the quantiles of the data (based on fitting the data into a normal distribution), 'defined' will use a predefined scale of 10 values specified by the user for each of the population_vulnerability_threshold values. If studying a single species use either 'interval' or 'quantile', if multiple species we recommend using the 'defined' option so that results across species datasets are comparable |
| population_vulnerability_thresholds | string of 10 variables separated by comma for defining the population_vulnerability of populations |
| vulnerability_rule | rule for calculating population vulnerability: # 1: Population vulnerability = mean score of Exposure, Neutral Sensitivity, Adaptive Sensitivity, Range shift potential, # 2: Population vulnerability = 70% weighting Exposure, 10% rest each, # 3: Population vulnerability = 70% weighting neutral sensitivity, 10% rest each, # 4: Population vulnerability = 70% weighting adaptive sensitivity, 10% rest each. # 5: Population vulnerability = 70% weighting range shift potential, 10% rest each |

**Table S2.** Summary of empirical genomic datasets, including links to raw genomic data.

| Species | Number of individuals | Publication | Data type | Sequencing | Data availability |
|---|---|---|---|---|---|
| *Myotis escalerai* | 216 | Razgour et al. 2019 | ddRAD-seq (SbfI, NlaIII) | 1 x Illumina HiSeq runs (162bp paired end reads) | PRJEB29086 |
| *Myotis crypticus* | 50 | Razgour et al. 2019 | ddRAD-seq (SbfI, NlaIII) | 1 x Illumina HiSeq runs (162bp paired end reads) | PRJEB29086 |
| *Afrixalus fornasini* | 43 | Barratt et al. 2018 | RAD-seq (*SbfI*) | 1 x Illumina HiSeq 2500 run (100 bp single end reads) | PRJNA472166 |

**Table S3.** Reclassified values assigned to map pixels for Globcover land cover data when preparing parameterised cumulative resistance surfaces for circuitscape analysis.

| Value | Globcover label | landcov1 | landcov2 | landcov3 |
|---|---|---|---|---|
| 11 | Post-flooding or irrigated croplands (or aquatic) | 50 | 50 | 100 |
| 14 | Rainfed croplands | 50 | 50 | 100 |
| 20 | Mosaic cropland (50-70%) / vegetation (grassland/shrubland/forest) (20-50%) | 10 | 10 | 10 |
| 30 | Mosaic vegetation (grassland/shrubland/forest) (50-70%) / cropland (20-50%) | 10 | 10 | 10 |
| 40 | Closed to open (>15%) broadleaved evergreen or semi-deciduous forest (>5m) | 5 | 5 | 5 |
| 50 | Closed (>40%) broadleaved deciduous forest (>5m) | 5 | 5 | 5 |
| 60 | Open (15-40%) broadleaved deciduous forest/woodland (>5m) | 5 | 5 | 5 |
| 70 | Closed (>40%) needleleaved evergreen forest (>5m) | 1 | 1 | 1 |
| 90 | Open (15-40%) needleleaved deciduous or evergreen forest (>5m) | 1 | 1 | 1 |
| 100 | Closed to open (>15%) mixed broadleaved and needleleaved forest (>5m) | 5 | 5 | 5 |
| 110 | Mosaic forest or shrubland (50-70%) / grassland (20-50%) | 20 | 20 | 20 |
| 120 | Mosaic grassland (50-70%) / forest or shrubland (20-50%) | 20 | 20 | 20 |
| 130 | Closed to open (>15%) (broadleaved or needleleaved, evergreen or deciduous) shrubland(<5m) | 40 | 50 | 100 |
| 140 | Closed to open (>15%) herbaceous vegetation (grassland, savannas or lichens/mosses) | 40 | 50 | 100 |
| 150 | Sparse (<15%) vegetation | 50 | 50 | 100 |
| 160 | Closed to open (>15%) broadleaved forest regularly flooded (semi-permanently or temporarily) - Fresh or brackish water | 50 | 50 | 100 |
| 170 | Closed (>40%) broadleaved forest or shrubland permanently flooded - Saline or brackish water | 50 | 50 | 100 |
| 180 | Closed to open (>15%) grassland or woody vegetation on regularly flooded or waterlogged soil - Fresh, brackish or saline water | 50 | 50 | 100 |
| 190 | Artificial surfaces and associated areas (Urban areas >50%) | 75 | 50 | 100 |
| 200 | Bare areas | 60 | 50 | 100 |
| 210 | Water bodies | NA | NA | NA |
| 220 | Permanent snow and ice | NA | NA | NA |
| 230 | No data (burnt areas, clouds,…) | NA | NA | NA |

**Table S4.** *Params.csv* file used to generate results for the 4 empirical datasets in this manuscript.

| Parameter | *Myotis crypticus* | *Myotis escalerai* | *Afrixalus fornasini* |
|---|---|---|---|
| species_binomial | Myotis_crypticus | Myotis_escalerai | Afrixalus_fornasini |
| user_specified_presences | no | no | no |
| gbif_limit | 1000 | 1000 | 1000 |
| gbif_date_limit_lower | 1900 | 1900 | 1900 |
| gbif_date_limit_upper | 2023 | 2023 | 2023 |
| climate_res | 2.5 | 2.5 | 2.5 |
| geographic_extent | -9.3,18.8,35,50.2 | -9.3,18.8,35,50.2 | 29.5,41.5,-21.5,-1.5 |
| current_climate_data_path | ./-data-/environmental_data/30s/current/ | ./-data-/environmental_data/30s/current/ | ./-data-/environmental_data/30s/current/ |
| future_climate_data_path | ./-data-/environmental_data/30s/future/HadGEM3-GC31-LL_ssp585_2061-2080/ | ./-data-/environmental_data/30s/future/HadGEM3-GC31-LL_ssp585_2061-2080/ | ./-data-B8/environmental_data/30s/future/HadGEM3-GC31-LL_ssp585_2061-2080/ |
| sp_rare_dist_km | 10 | 10 | 10 |
| sdm_reps_per_algorithm | 10 | 10 | 10 |
| bg_buff_dist_degrees | 2 | 2 | 2 |
| n_background_points | 10000 | 10000 | 10000 |
| perform_vif | no | no | yes |
| subset_predictors | bioclim_1,bioclim_4,bioclim_7,bioclim_5,bioclim_6,slope,land_cover | bioclim_1,bioclim_4,bioclim_7,bioclim_5,bioclim_6,slope,land_cover | |
| biomod_algorithms | CTA','MAXENT.Phillips' | CTA','MAXENT.Phillips' | RF','GAM','MAXENT.Phillips' |
| maxent_path | ./-data-/ | ./-data-/ | ./-data-/ |
| data_split_percentage | 80 | 80 | 80 |
| ROC_min | 0.75 | 0.75 | 0.75 |
| TSS_min | | | |

| | | | |
|---|---|---|---|
| env_predictor_1 | bioclim_5 | bioclim_5 | bioclim_5 |
| env_predictor_2 | bioclim_18 | bioclim_18 | bioclim_18 |
| k_min | 2 | 2 | 2 |
| k_max | 10 | 10 | 10 |
| snmf_reps | 10 | 10 | 10 |
| ploidy | 2 | 2 | 2 |
| plink_executable | ./-data-/plink | ./-data-/plink | ./-data-/plink |
| k | 3 | 3 | 4 |
| scale_gif_lfmm | 0.63 | 0.65 | 0.75 |
| rda_axes | 2 | 2 | 2 |
| rda_SD | 3 | 3 | 3 |
| scale_gif_rda | 0.9 | 0.9 | 0.9 |
| which_loci | 6 | 6 | 1 |
| category_1 | hot_dry | hot_dry | hot_dry |
| category_2 | cold_wet | cold_wet | cold_wet |
| circuitscape_transform_zeros | yes | yes | yes |
| circuitscape_layers | current_sdm_resistance,slope_resistance,landcov1_resistance,env_1_resistance,env_2_resistance | current_sdm_resistance,slope_resistance,landcov1_resistance,env_1_resistance,env_2_resistance | current_sdm_resistance,slope_resistance,landcov1_resistance,env_1_resistance,env_2_resistance |
| circuitscape_weights | 0.25,0.1,0.25,0.2,0.2 | 0.25,0.1,0.25,0.2,0.2 | 0.25,0.1,0.25,0.2,0.2 |
| max_dispersal_distance_km | 464 | 464 | 100 |
| exposure_rule | 1 | 1 | 1 |
| vulnerability_rule | 1 | 1 | 1 |
| exposure_quantification | interval | interval | interval |
| exposure_env_1_thresholds | 3,3.5,4,4.5,5,5.5,6,6.5,7,7.5 | 3,3.5,4,4.5,5,5.5,6,6.5,7,7.5 | 3,3.5,4,4.5,5,5.5,6,6.5,7,7.5 |
| exposure_env_2_thresholds | -100,-50,0,50,100,150,200,250,300,350 | -100,-50,0,50,100,150,200,250,300,350 | -100,-50,0,50,100,150,200,250,300,350 |
| exposure_sdm_thresholds | -90,-80,-70,-60,-50,-40,-30,-20,-10,0 | -90,-80,-70,-60,-50,-40,-30,-20,-10,0 | -90,-80,-70,-60,-50,-40,-30,-20,-10,0 |

| | | | |
|---|---|---|---|
| neutral_sensitivity_quantification | interval | interval | interval |
| neutral_sensitivity_thresholds | 0,0.1,0.2,0.3,0.4,0.5,0.6,0.7,0.8,0.9 | 0,0.1,0.2,0.3,0.4,0.5,0.6,0.7,0.8,0.9 | 0,0.1,0.2,0.3,0.4,0.5,0.6,0.7,0.8,0.9 |
| adaptive_sensitivity_quantification | interval | interval | interval |
| adaptive_sensitivity_thresholds | 0,0.1,0.2,0.3,0.4,0.5,0.6,0.7,0.8,0.9 | 0,0.1,0.2,0.3,0.4,0.5,0.6,0.7,0.8,0.9 | 0,0.1,0.2,0.3,0.4,0.5,0.6,0.7,0.8,0.9 |
| range_shift_potential_quantification | interval | interval | interval |
| range_shift_potential_thresholds | 0,0.1,0.2,0.3,0.4,0.5,0.6,0.7,0.8,0.9 | 0,0.1,0.2,0.3,0.4,0.5,0.6,0.7,0.8,0.9 | 0,0.1,0.2,0.3,0.4,0.5,0.6,0.7,0.8,0.9 |
| pop_vulnerability_quantification | interval | interval | interval |
| pop_vulnerability_thresholds | 1,2,3,4,5,6,7,8,9,10 | 1,2,3,4,5,6,7,8,9,10 | 1,2,3,4,5,6,7,8,9,10 |

### **Appendix S1.** Complete log file for *Afrixalus fornasini.*

BEGIN...
Life on the edge pipeline started for Afrixalus_fornasini  at 2023-05-26 17:08:59 CEST

--------------------------------------------------------------
Preparing spatial data...
--------------------------------------------------------------
> Preparing spatial presence data...
> Spatial presence data downloaded and prepared successfully...
> Number of georeferenced genomic samples:  43
> Number of GBIF records downloaded:  298
> Output presence data saved

> Preparing environmental predictor data...
> Environmental predictor data downloaded and prepared successfully...
> Current Worldclim2 climate data prepared at 0.008333333 km^2 resolution
> Future Worldclim2 climate data prepared at  0.008333333  km^2 resolution
> Output raster data saved

> Extracting predictor data for each sample (for later environmental dissimilarity and GEA analyses)...
> DONE...
> Spatially rarefying presence data...
> Spatial rarefication of presence data completed sucessfully...
> Rarefied the presence data to retain only presences a minimum distance of 10 km from nearest sample
> Total number of presences before spatial rarefication: 341
> Total number of presences after spatial rarefication: 113
> Output spatially rarefied presence data saved and plotted

> Generating and plotting random background (pseudoabsence) points for species distribution modelling...
> Number of cells in modelling area:  272448
> Number of cells in background buffer ( 2  degrees):  68314
> Generating background points from this region (n= 10000 )
> Background (pseudoabsence) data prepared successfully...
>  10000  background points generated in a buffer radius of 2 degrees around presence points
> Output background points saved and plotted

--------------------------------------------------------------
Species Distribution Modelling using biomod2 package...
--------------------------------------------------------------
> Building SDMs for SDM algorithms -  RF','GAM','MAXENT.Phillips' , using 10 replicates per algorithm
> Number of background points:  10000> Building final ensemble model with passed thresholds ROC> 0.75 ...
>>> Output SDMs (present and future) saved and plotted
> SDMs completed...

--------------------------------------------------------------

```
Calculating environmental dissimilarity and quantifying exposure...
-------------------------------------------------------------
> Reading SDMs and predictor variables for present and future (2070)...
> Environmental dissimilarity of SDMs, env predictors, plots and rasters saved for present-future...
> Environmental dissimilarity  completed ...

> Now calculating Exposure based on magnitude of change predicted at each location...
Exposure calculated based on the mean score for SDM, bioclim_5 , and bioclim_18 ...
> Exposure calculations completed successfully ...

-------------------------------------------------------------
Assessing population structure and imputing missing data...
-------------------------------------------------------------
> Running sNMF population structure analysis...
> Range of K tested: 2 : 10
> K with lowest cross-entropy: 2
> Output sNMF population assignment data saved and plotted
> sNMF population structure analysis completed successfully...

> Imputing missing data. using LEA..
> Imputed data saved, LEA completed successfully...

> Imputing missing data using mean genotypes..
> Imputed data saved, mean genotype imputation completed successfully...

> Running PCA population structure analysis...
> Output PCA population assignment data saved and plotted

** WARNING: check this plot to identify number of population clusters before going further
> PCA  population structure analysis completed successfully...

-------------------------------------------------------------
Running LFMM (Latent Factor Mixed Models)...
-------------------------------------------------------------
> Assessing correlations of predictor variables...
> Correlations of predictor variables checked and completed...
> Variance Inflation Factors for variables are:  1.060546 1.060546
>>> Output correlation data saved and plotted

> Running LFMM (Latent Factor Mixed Models)...
** WARNING: Decision needed for number of populations...

> Evaluate sNMF and PCA population structure outputs to define K
> Predictor variable bioclim_5 ...
> Calculated GIF for predictor variable bioclim_5  is:  0.9643844
** WARNING: Examine the unadjusted vs. adjusted p-values, if adjustment is too conservative (distribution is too flat, increase the GIF...)
> Updated GIF is:  0.7232883  (original GIF* 0.75 )
>>> Output summary of adjusted vs. unadjusted p-values plotted
>>> Output Manhattan plot of adjusted p-values saved...
> Predictor variable bioclim_18 ...
> Calculated GIF for predictor variable bioclim_18  is:  1.141314
** WARNING: Examine the unadjusted vs. adjusted p-values, if adjustment is too conservative (distribution is too flat, increase the GIF...)
> Updated GIF is:  0.8559853  (original GIF* 0.75 )
>>> Output summary of adjusted vs. unadjusted p-values plotted
>>> Output Manhattan plot of adjusted p-values saved...
>>> Output candidate SNPs by predictor saved
>>> Output candidate SNPs (merged) saved
>>> Candidate SNPs FDR<0.1: 726
>>> Candidate SNPs FDR<0.05: 562
>>> Candidate SNPs FDR<0.01: 135
> LFMM analysis completed successfully...

-------------------------------------------------------------
Running RDA (Redundancy Analysis)...
-------------------------------------------------------------
** WARNING: Decision needed - how many axes to retain for the RDA? Default is to match the number of predictors - in this case:  2 ( bioclim_5 ,
bioclim_18 )
> Output candidate SNPs saved and plotted
> Calculated GIF for predictor variables is:  1.398412
> Updated GIF is:  1.258571  (original GIF* 0.9 )
```

> Output summary of adjusted vs. unadjusted p-values plotted
> Output plot of candidate SNPs (post cut-off) saved and plotted
> Candidate SNPs SD± 3 : 12
> Candidate SNPs FDR<0.1: 0
> Candidate SNPs FDR<0.05: 0
> Candidate SNPs FDR<0.01: 0
> RDA analysis completed successfully...

--------------------------------------------------------------
Categorising individuals (using RDA)...
--------------------------------------------------------------
Reading in environmental and genomic data...
Subsetting adaptive loci present in either LFMM (FDR<0.05) or RDA (SD<3) analyses...
Number of loci:  574
Performing RDA to categorise individuals...
Writing output plots...
> Categorisation of individuals using RDA completed successfully...

--------------------------------------------------------------
Quantifying adaptive diversity ...
--------------------------------------------------------------
> Reading categorised individuals based on step 9a...
> Summarising samples at each unique lat/long coords ...
> Plotting categorised individuals ...
> DONE ...
> Plotting categorised populations ... (all unique lat/long coords with > 1 sample)...
> DONE ...
> Plotting individual categorisation maps...
> Output plots and data written...

> Adaptive sensitivity analyses completed successfully...

--------------------------------------------------------------
Quantifying neutral diversity ...
--------------------------------------------------------------
> Running PLINK to calculate heterozygosity (including adaptive regions)...
> Heterozygosity calculated...
> Now masking out adaptive regions and recalculating neutral heterozygosity...
> Neutral heterozygosity calculated...
> Summarising outputs based on unique lat/long coordinates...
> Creating neutral sensitivity metrics per population...
> Neutral sensitivity analyses completed successfully ...

--------------------------------------------------------------
Calculating Sensitivity...
--------------------------------------------------------------
> Generating Sensitivity metrics...
> hot_dry sensitivity calculated
> cold_wet sensitivity calculated
> adaptive sensitivity calculated based on hot_dry and cold_wet individuals
> Sensitivity analyses completed successfully...

--------------------------------------------------------------
12. Creating Circuitscape *.ini and *.jl files...
--------------------------------------------------------------
> Creating points file from georeferenced genomic samples... DONE!
> Creating Circuitscape *.ini file...> DONE!
> Creating Circuitscape *.jl file...> DONE!
> DONE!
> Circuitscape files created successfully...

--------------------------------------------------------------
14. Calculating Range shift potential for each population based on Circuitscape outputs...
--------------------------------------------------------------
> Plotted Circuitscape landscape connectivity map...
> Calculated range shift potential metrics and wrote to file...
> Range shift potential analyses completed...

--------------------------------------------------------------
Summarising analyses and generating final population vulnerability scores...

--------------------------------------------------------------
Population vulnerability calculated based on mean of Exposure, Neutral Sensitivity, Adaptive Sensitivity, and Range shift Potential...
Exposure scores calculated, saved and plotted...
Now interpolating the data...
Exposure scores interpolated and plotted...

Neutral sensitivity scores calculated, saved and plotted...
Now interpolating the data...
Neutral sensitivity scores interpolated and plotted...

Adaptive sensitivity scores plotted...
Now interpolating the data...
Adaptive sensitivity scores interpolated and plotted...

Range shift potential scores saved and plotted...
Now interpolating the data...
Range shift potential scores interpolated and plotted...

Plotting composite panels of Exposure, Neutral/Adaptive diversity, Range shift potential...

Population vulnerability scores calculated, saved and plotted...
Now interpolating the data...
Final Population vulnerability scores interpolated and plotted...
Population vulnerability completed successfully...

--------------------------------------------------------------------------------
Generating final PDFs with all summary statistics, plots and information...
--------------------------------------------------------------------------------
Summarising relevant outputs to create summary PDFs...
DONE!
END...

Life on the edge pipeline completed for Afrixalus_fornasini at 2023-05-27 02:04:24 CEST

**Appendix S2**. Final summary PDF for *Afrixalus fornasini (overleaf)*.
