## Supporting Information 2 for "Life on the edge: a new toolbox for population-level climate change vulnerability assessments"

SPATIAL DATA

> Number of georeferenced genomic samples:  43

> Number of GBIF records downloaded:  298

> Total number of presences before spatial rarefication: 341

> Total number of presences after spatial rarefication: 113

Number of bg points in SDMs: 10000

Geographic extent (xmin,xmax, ymin,ymax): 29.5,41.5,−21.5,−1.5

> Resolution: 0.008333333 km^2 resolution

SDMs

SDM replicates per algorithm: 10

Sampling strategy:

Data split: training/testing: 80%

Min AUC threshold for model to be included in final ensemble: 0.75

Min TSS threshold for model to be included in final ensemble: 0.75

Models tested: RF','GAM','MAXENT.Phillips'

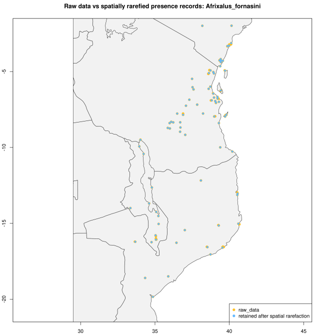


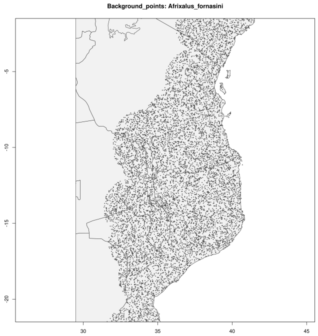

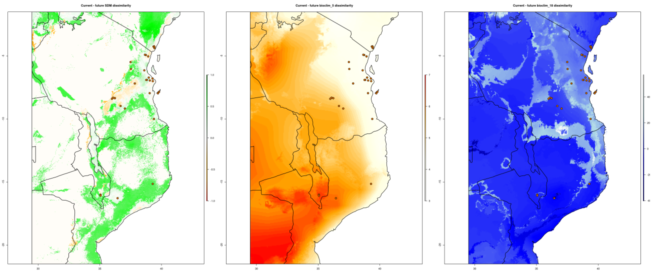
Current - future SDM dissimilarity
Current - future bioclim_5 dissimilarity
Current - future bioclim_18 dissimilarity

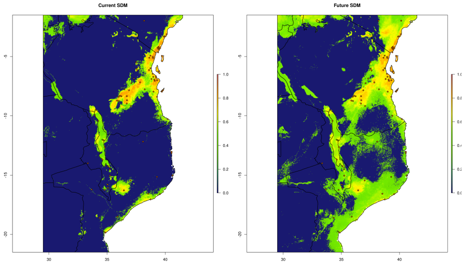
Current SDM
Future SDM

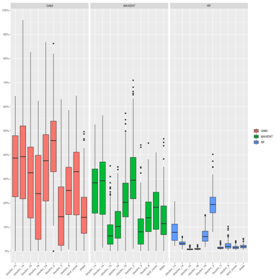
GAM
MAXENT
RF

| Pop | SDM exposure | env1 exposure | env2 exposure | exposure |
|---|---|---|---|---|
| 1 | 6 | 10 | 2 | 6 |
| 2 | 7 | 1 | 10 | 6 |
| 3 | 5 | 1 | 7 | 4.33 |
| 4 | 7 | 1 | 6 | 4.67 |
| 5 | 8 | 6 | 3 | 5.67 |
| 6 | 6 | 1 | 6 | 4.33 |
| 7 | 7 | 1 | 8 | 5.33 |
| 8 | 7 | 1 | 8 | 5.33 |
| 9 | 7 | 1 | 6 | 4.67 |
| 10 | 8 | 1 | 5 | 4.67 |
| 11 | 6 | 1 | 5 | 4 |
| 12 | 7 | 6 | 3 | 5.33 |
| 13 | 8 | 1 | 9 | 6 |
| 14 | 7 | 1 | 4 | 4 |
| 15 | 6 | 1 | 6 | 4.33 |
| 16 | 5 | 1 | 5 | 3.67 |
| 17 | 1 | 1 | 5 | 2.33 |
| 18 | 6 | 1 | 6 | 4.33 |
| 19 | 6 | 1 | 8 | 5 |
| 20 | 7 | 6 | 9 | 7.33 |
| 21 | 8 | 5 | 3 | 5.33 |
| 22 | 8 | 1 | 4 | 4.33 |
| 23 | 6 | 1 | 4 | 3.67 |
| 24 | 7 | 1 | 4 | 4 |
| 25 | 6 | 1 | 4 | 3.67 |
| 26 | 3 | 5 | 1 | 3 |
| 27 | 6 | 9 | 1 | 5.33 |
| 28 | 10 | 10 | 2 | 7.33 |
| 29 | 10 | 6 | 3 | 6.33 |
| 30 | 10 | 1 | 6 | 5.67 |
| 31 | 8 | 1 | 6 | 5 |
| 32 | 7 | 6 | 3 | 5.33 |

Exposure - interpolated

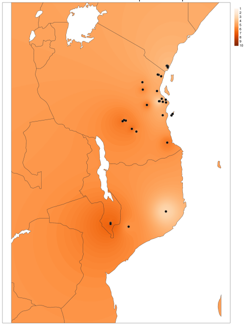

ADAPTIVE LOCI

Number of LFMM candidates FDR<0.1: 726

Number of LFMM candidates FDR<0.05: 562

Number of LFMM candidates FDR<0.01: 135

Number of RDA candidates loci FDR<0.1: 0

Number of RDA candidates FDR<0.05: 0

Number of RDA candidates FDR<0.01: 0

Number of RDA candidates SD<3: 12

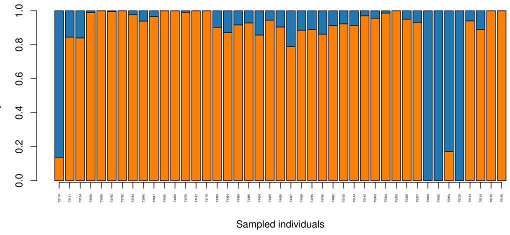


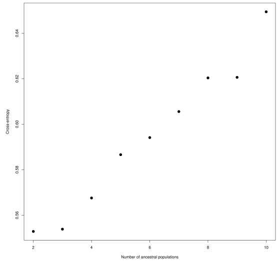


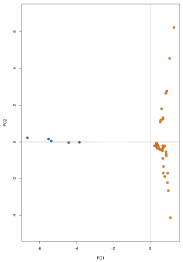


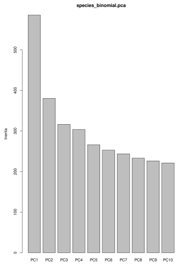

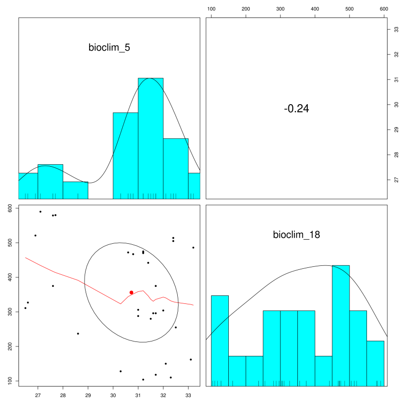
bioclim_5
-0.24
bioclim_18

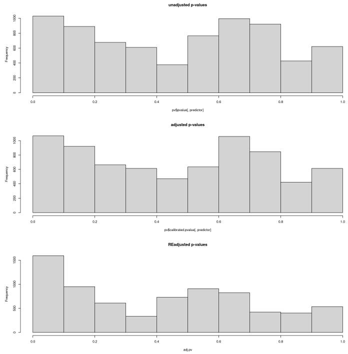
unadjusted p-values
pv$pvalue[, predictor]
adjusted p-values
pv$calibrated.pvalue[, predictor]
REadjusted p-values
adj.pv
Frequency

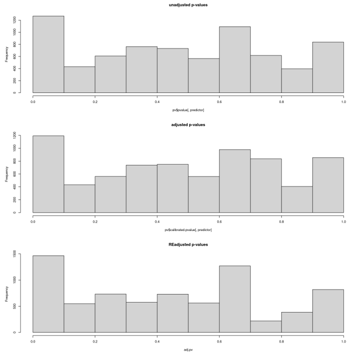
unadjusted p-values
pv$pvalue[, predictor]
adjusted p-values
pv$calibrated.pvalue[, predictor]
REadjusted p-values
adj.pv
Frequency

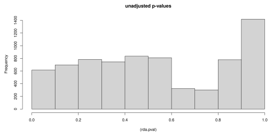
unadjusted p-values
Frequency
(rda.pval)

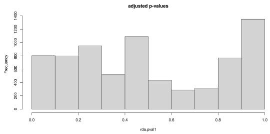
adjusted p-values
Frequency
rda.pval1

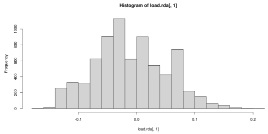
Histogram of load.rda[, 1]
Frequency
load.rda[, 1]

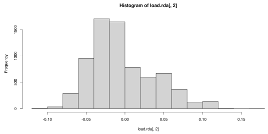
Histogram of load.rda[, 2]
Frequency
load.rda[, 2]

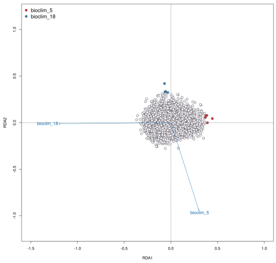
bioclim_5
bioclim_18
RDA2
RDA1

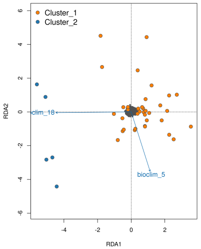
Cluster_1
Cluster_2
clim_18
bioclim_5
RDA2
RDA1

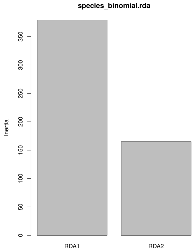
species_binomial.rda
Inertia
RDA1
RDA2

LOCALLY ADAPTED INDIVIDUALS

Number of hot_dry adapted individuals: 16

Number of cold_wet adapted individuals:

10

Number of intermediate (i.e. not locally

adapted) individuals: 17

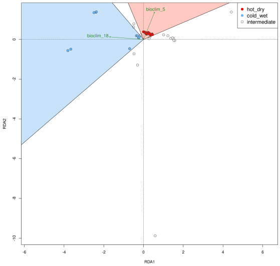


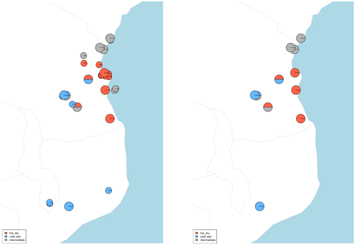

| Pop | Neutral observed heterozygosity | Neutral sensitivity |
|---|---|---|
| 1 | 0.18 | 8 |
| 2 | 0.13 | 8 |
| 3 | 0.52 | 2 |
| 4 | 0.39 | 4 |
| 5 | 0.36 | 5 |
| 6 | 0.29 | 6 |
| 7 | 0.45 | 3 |
| 8 | 0.46 | 3 |
| 9 | 0.33 | 5 |
| 10 | 0.35 | 5 |
| 11 | 0.14 | 8 |
| 12 | 0.11 | 9 |
| 13 | 0.19 | 7 |
| 14 | 0.27 | 6 |
| 15 | 0.21 | 7 |
| 16 | 0.15 | 8 |
| 17 | 0.19 | 7 |
| 18 | 0.13 | 8 |
| 19 | 0.13 | 8 |
| 20 | 0.19 | 7 |
| 21 | 0.21 | 7 |
| 22 | 0.25 | 6 |
| 23 | 0.4 | 4 |
| 24 | 0.17 | 8 |
| 25 | 0.12 | 9 |
| 26 | 0.35 | 5 |
| 27 | 0.22 | 7 |
| 28 | 0.32 | 5 |
| 29 | 0.36 | 5 |
| 30 | 0 | 10 |
| 31 | 0.34 | 5 |
| 32 | 0.61 | 1 |

Neutral sensitivity - interpolated

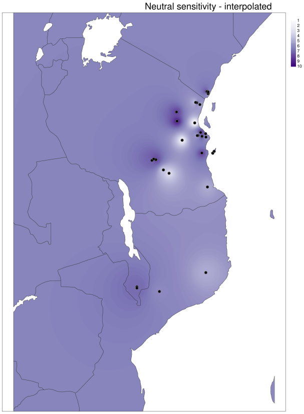

| pop | hot_dry | cold_wet | intermediate | category_1_sensitivity | category_2_sensitivity | adaptive_sensitivity | adaptive_sensitivity_metric |
|---|---|---|---|---|---|---|---|
| 1 | 0 | 2 | 0 | 0 | −0.16 | −0.16 | 1 |
| 2 | 0 | 1 | 0 | 0 | −0.03 | −0.03 | 3 |
| 3 | 0 | 1 | 0 | 0 | −0.02 | −0.02 | 3 |
| 4 | 0 | 1 | 0 | 0 | −0.1 | −0.1 | 2 |
| 5 | 2 | 0 | 0 | 0.12 | 0 | 0.12 | 7 |
| 6 | 1 | 0 | 1 | 0.08 | 0 | 0.08 | 6 |
| 7 | 0 | 1 | 0 | 0 | 0 | 0 | 4 |
| 8 | 0 | 0 | 1 | 0 | 0 | 0 | 4 |
| 9 | 0 | 0 | 1 | 0 | 0 | 0 | 4 |
| 10 | 0 | 1 | 1 | 0 | 0.01 | 0.01 | 4 |
| 11 | 0 | 2 | 0 | 0 | 0.01 | 0.01 | 4 |
| 12 | 0 | 0 | 1 | 0 | 0 | 0 | 4 |
| 13 | 2 | 0 | 0 | 0.12 | 0 | 0.12 | 7 |
| 14 | 1 | 0 | 0 | 0.12 | 0 | 0.12 | 7 |
| 15 | 0 | 0 | 1 | 0 | 0 | 0 | 4 |
| 16 | 1 | 1 | 0 | 0.06 | 0.23 | 0.29 | 10 |
| 17 | 1 | 0 | 0 | 0.12 | 0 | 0.12 | 7 |
| 18 | 1 | 0 | 0 | 0.12 | 0 | 0.12 | 7 |
| 19 | 1 | 0 | 0 | 0.11 | 0 | 0.11 | 7 |
| 20 | 1 | 0 | 0 | 0.11 | 0 | 0.11 | 7 |
| 21 | 1 | 0 | 0 | 0.11 | 0 | 0.11 | 7 |
| 22 | 2 | 0 | 0 | 0.11 | 0 | 0.11 | 7 |
| 23 | 1 | 0 | 0 | 0.11 | 0 | 0.11 | 7 |
| 24 | 1 | 0 | 0 | 0.13 | 0 | 0.13 | 7 |
| 25 | 0 | 0 | 1 | 0 | 0 | 0 | 4 |
| 26 | 0 | 0 | 2 | 0 | 0 | 0 | 4 |
| 27 | 0 | 0 | 1 | 0 | 0 | 0 | 4 |
| 28 | 0 | 0 | 2 | 0 | 0 | 0 | 4 |
| 29 | 0 | 0 | 1 | 0 | 0 | 0 | 4 |
| 30 | 0 | 0 | 1 | 0 | 0 | 0 | 4 |
| 31 | 0 | 0 | 1 | 0 | 0 | 0 | 4 |
| 32 | 0 | 0 | 2 | 0 | 0 | 0 | 4 |

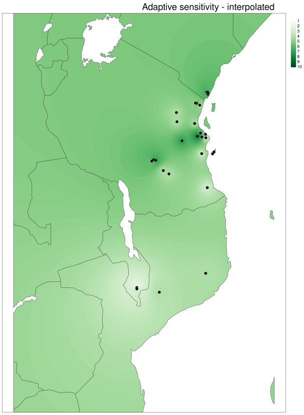

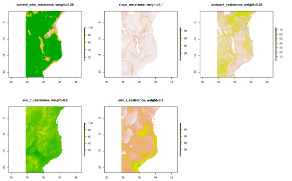


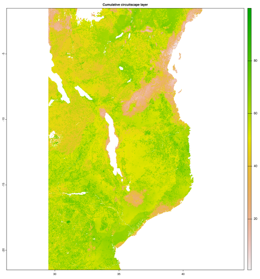


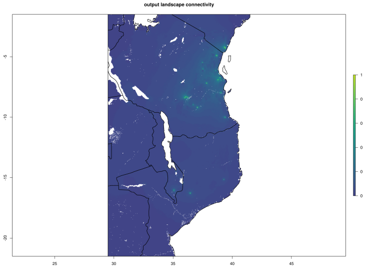


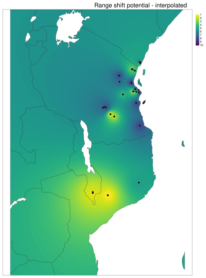


| Pop | mean_connectivity_normalised | range shift potential |
|---|---|---|
| 1 | 0.99 | 2 |
| 2 | 0.81 | 8 |
| 3 | 0.81 | 8 |
| 4 | 0.42 | 10 |
| 5 | 0.65 | 9 |
| 6 | 0.42 | 10 |
| 7 | 0.42 | 10 |
| 8 | 0 | 10 |
| 9 | 0.72 | 9 |
| 10 | 0.98 | 4 |
| 11 | 0.98 | 4 |
| 12 | 0.99 | 2 |
| 13 | 0.98 | 4 |
| 14 | 0.98 | 3 |
| 15 | 0.98 | 3 |
| 16 | 0.98 | 3 |
| 17 | 0.99 | 1 |
| 18 | 0.99 | 1 |
| 19 | 0.87 | 5 |
| 20 | 0.87 | 5 |
| 21 | 0.85 | 7 |
| 22 | 0.87 | 5 |
| 23 | 0.99 | 2 |
| 24 | 0.85 | 7 |
| 25 | 0.87 | 5 |
| 26 | 0.87 | 5 |
| 27 | 0.99 | 1 |
| 28 | 0.99 | 1 |
| 29 | 1 | 1 |
| 30 | 0.84 | 7 |
| 31 | 0.81 | 8 |
| 32 | 0.81 | 8 |

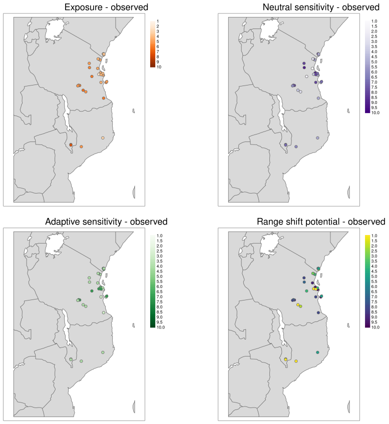
Exposure - observed
Neutral sensitivity - observed
Adaptive sensitivity - observed
Range shift potential - observed

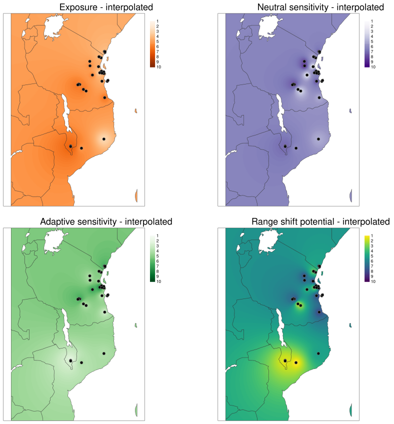
Exposure - interpolated
Neutral sensitivity - interpolated
Adaptive sensitivity - interpolated
Range shift potential - interpolated

| Pop | Exposure | Neutral_sensitivity | Adaptive_sensitivity | Range shift potential | Population vulnerability |
|---|---|---|---|---|---|
| 1 | 6 | 7 | 1 | 2 | 4 |
| 2 | 6 | 5 | 3 | 8 | 5.5 |
| 3 | 4.33 | 8 | 3 | 8 | 5.83 |
| 4 | 4.67 | 5 | 2 | 10 | 5.42 |
| 5 | 5.67 | 8 | 7 | 9 | 7.42 |
| 6 | 4.33 | 9 | 6 | 10 | 7.33 |
| 7 | 5.33 | 5 | 4 | 10 | 6.08 |
| 8 | 5.33 | 7 | 4 | 10 | 6.58 |
| 9 | 4.67 | 1 | 4 | 9 | 4.67 |
| 10 | 4.67 | 5 | 4 | 4 | 4.42 |
| 11 | 4 | 7 | 4 | 4 | 4.75 |
| 12 | 5.33 | 3 | 4 | 2 | 3.58 |
| 13 | 6 | 2 | 7 | 4 | 4.75 |
| 14 | 4 | 8 | 7 | 3 | 5.5 |
| 15 | 4.33 | 3 | 4 | 3 | 3.58 |
| 16 | 3.67 | 7 | 10 | 3 | 5.92 |
| 17 | 2.33 | 4 | 7 | 1 | 3.58 |
| 18 | 4.33 | 8 | 7 | 1 | 5.08 |
| 19 | 5 | 7 | 7 | 5 | 6 |
| 20 | 7.33 | 8 | 7 | 5 | 6.83 |
| 21 | 5.33 | 8 | 7 | 7 | 6.83 |
| 22 | 4.33 | 6 | 7 | 5 | 5.58 |
| 23 | 3.67 | 5 | 7 | 2 | 4.42 |
| 24 | 4 | 10 | 7 | 7 | 7 |
| 25 | 3.67 | 5 | 4 | 5 | 4.42 |
| 26 | 3 | 5 | 4 | 5 | 4.25 |
| 27 | 5.33 | 6 | 4 | 1 | 4.08 |
| 28 | 7.33 | 7 | 4 | 1 | 4.83 |
| 29 | 6.33 | 4 | 4 | 1 | 3.83 |
| 30 | 5.67 | 9 | 4 | 7 | 6.42 |
| 31 | 5 | 8 | 4 | 8 | 6.25 |
| 32 | 5.33 | 6 | 4 | 8 | 5.83 |

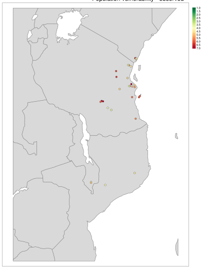
Population vulnerability - observed

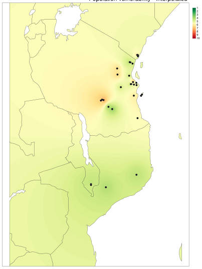
Population vulnerability - interpolated
