## Supporting Information 3 for "Life on the edge: a new toolbox for population-level climate change vulnerability assessments"

### **Appendix S3.** Complete log file for *Myotis escalerai.*

BEGIN...
Life on the edge pipeline started for Myotis_escalerai at 2023-05-13 11:15:06 CET

--------------------------------------------------------------
Preparing spatial data...
--------------------------------------------------------------
> Preparing spatial presence data...
> Spatial presence data downloaded and prepared successfully...
> Number of GBIF records downloaded: 300
> Number of georeferenced genomic samples: 216
>>> Output presence data saved

> Preparing environmental predictor data...
> Environmental predictor data downloaded and prepared successfully...
> Current Worldclim2 climate data prepared at 0.008333333 km^2 resolution
> Future Worldclim2 climate data prepared at 0.008333333 km^2 resolution
>>> Output raster data saved

> Extracting predictor data for each sample (for later environmental dissimilarity and GEA analyses)...
> DONE...
> Spatially rarefying presence data...
> Spatial rarefication of presence data completed sucessfully...
> Rarefied the presence data to retain only presences a minimum distance of 10 km from nearest sample
> Total number of presences before spatial rarefication: 516
> Total number of presences after spatial rarefication: 122
>>> Output spatially rarefied presence data saved and plotted

> Generating and plotting random background (pseudoabsence) points for species distribution modelling...
> Number of cells in modelling area: 343998
> Number of cells in background buffer ( 2 degrees): 42157
> Generating background points from this region (n= 10000 )
> Background (pseudoabsence) data prepared successfully...
> 10000 background points generated in a buffer radius of 2 degrees around presence points
>>> Output background points saved and plotted

--------------------------------------------------------------
Species Distribution Modelling using biomod2 package...
--------------------------------------------------------------
> Predictor variables checked for autocorrelation using sample locations and Variance Inflation Factors...
> Predictors remaining:
class : RasterStack dimensions : 1824, 3372, 6150528, 6 (nrow, ncol, ncell, nlayers) resolution : 0.008333333, 0.008333333 (x, y) extent : -9.3, 18.8, 35, 50.2 (xmin, xmax, ymin, ymax) crs : +proj=longlat +datum=WGS84 +no_defs names : bioclim_1, bioclim_4, bioclim_7, bioclim_5, bioclim_6, elevation min values : -11.09583, 246.90353, 10.10000, -1.70000, -18.90000, -57.00000 max values : 20.57083, 853.91998, 37.00000, 40.20000, 11.70000, 4482.00000
> Building SDMs for SDM algorithms - CTA','MAXENT.Phillips' , using 10 replicates per algorithm
> Number of background points: 10000> Building final ensemble model with passed thresholds ROC> 0.75 ...
>>> Output SDMs (present and future) saved and plotted
> SDMs completed...

--------------------------------------------------------------
Assessing population structure and imputing missing data...
--------------------------------------------------------------
> Running sNMF population structure analysis...
> Range of K tested: 2 : 10
> K with lowest cross-entropy: 4
>>> Output sNMF population assignment data saved and plotted
> sNMF population structure analysis completed successfully...

> Imputing missing data. using LEA..
> Imputed data saved, LEA completed successfully...

> Imputing missing data using mean genotypes..
> Imputed data saved, mean genotype imputation completed successfully...

> Running PCA population structure analysis...
>>> Output PCA population assignment data saved and plotted

** WARNING: check this plot to identify number of population clusters before going further

```
> PCA  population structure analysis completed successfully...

--------------------------------------------------------------
Running LFMM (Latent Factor Mixed Models)...
--------------------------------------------------------------
> Assessing correlations of predictor variables...
> Correlations of predictor variables checked and completed...
> Variance Inflation Factors for variables are:  2.769715 2.769715
>>> Output correlation data saved and plotted

> Running LFMM (Latent Factor Mixed Models)...
** WARNING: Decision needed for number of populations...

> Evaluate sNMF and PCA population structure outputs to define K
> Predictor variable bioclim_5 ...
> Calculated GIF for predictor variable bioclim_5  is:  1.449053
** WARNING: Examine the unadjusted vs. adjusted p-values, if adjustment is too conservative (distribution is too flat, increase the GIF...)
> Updated GIF is:  0.9418842  (original GIF* 0.65 )
>>> Output summary of adjusted vs. unadjusted p-values plotted
>>> Output Manhattan plot of adjusted p-values saved...
> Predictor variable bioclim_18 ...
> Calculated GIF for predictor variable bioclim_18  is:  1.81233
** WARNING: Examine the unadjusted vs. adjusted p-values, if adjustment is too conservative (distribution is too flat, increase the GIF...)
> Updated GIF is:  1.178015  (original GIF* 0.65 )
>>> Output summary of adjusted vs. unadjusted p-values plotted
>>> Output Manhattan plot of adjusted p-values saved...
>>> Output candidate SNPs by predictor saved
>>> Output candidate SNPs (merged) saved
> LFMM analysis completed successfully...

--------------------------------------------------------------
Running RDA (Redundancy Analysis)...
--------------------------------------------------------------
** WARNING: Decision needed - how many axes to retain for the RDA? Default is to match the number of predictors - in this case:  2 ( bioclim_5 , bioclim_18 )
>>> Output candidate SNPs saved and plotted
> Calculated GIF for predictor variables is:  1.337985
> Updated GIF is:  1.204186  (original GIF* 0.9 )
>>> Output summary of adjusted vs. unadjusted p-values plotted
>>> Output plot of candidate SNPs (post cut-off) saved and plotted
> RDA analysis completed successfully...

--------------------------------------------------------------
Categorising individuals (using RDA)...
--------------------------------------------------------------
Reading in environmental and genomic data...
Subsetting adaptive loci present in both LFMM (FDR<0.05) and RDA (SD<3) analyses...
Number of loci:  50
Performing RDA to categorise individuals...
Writing output plots...
> Categorisation of individuals using RDA completed successfully...

--------------------------------------------------------------
Quantifying adaptive diversity ...
--------------------------------------------------------------
> Reading categorised individuals based on step 9a...
> Summarising samples at each unique lat/long coords ...
> Plotting categorised individuals ...
> DONE ...
> Plotting categorised populations ... (all unique lat/long coords with > 1 sample)...
> DONE ...
> Plotting individual categorisation maps...
> Output plots and data written...

> Adaptive sensitivity analyses completed successfully...
```

#### **Appendix S4.** Complete log file for *Myotis crypticus*.

BEGIN...
Life on the edge pipeline started for Myotis_crypticus  at 2023-05-13 11:15:16 CET

--------------------------------------------------------------
Preparing spatial data...
--------------------------------------------------------------
> Preparing spatial presence data...
> Spatial presence data downloaded and prepared successfully...
> Number of GBIF records downloaded:  198
> Number of georeferenced genomic samples:  50
>>> Output presence data saved

> Preparing environmental predictor data...
> Environmental predictor data downloaded and prepared successfully...
> Current Worldclim2 climate data prepared at 0.008333333 km^2 resolution
> Future Worldclim2 climate data prepared at  0.008333333  km^2 resolution
>>> Output raster data saved

> Extracting predictor data for each sample (for later environmental dissimilarity and GEA analyses)...
> DONE...
> Spatially rarefying presence data...
> Spatial rarefication of presence data completed sucessfully...
> Rarefied the presence data to retain only presences a minimum distance of 10 km from nearest sample
> Total number of presences before spatial rarefication: 248
> Total number of presences after spatial rarefication: 89
>>> Output spatially rarefied presence data saved and plotted

> Generating and plotting random background (pseudoabsence) points for species distribution modelling...
> Number of cells in modelling area:  122958
> Number of cells in background buffer ( 2  degrees):  50949
> Generating background points from this region (n= 10000 )
> Background (pseudoabsence) data prepared successfully...
>  10000  background points generated in a buffer radius of 2 degrees around presence points
>>> Output background points saved and plotted

--------------------------------------------------------------
Species Distribution Modelling using biomod2 package...
--------------------------------------------------------------
> Predictor variables checked for autocorrelation using sample locations and Variance Inflation Factors...
> Predictors remaining:
class          : RasterStack  dimensions : 1824, 3372, 6150528, 6  (nrow, ncol, ncell, nlayers) resolution : 0.008333333, 0.008333333  (x, y) extent       : -9.3, 18.8, 35, 50.2  (xmin, xmax, ymin, ymax) crs          : +proj=longlat +datum=WGS84 +no_defs  names          :  bioclim_1, bioclim_4,  bioclim_7,  bioclim_5,  bioclim_6,  elevation  min values :  -11.09583,  246.90353,   10.10000,   -1.70000,  -18.90000,  -57.00000  max values :   20.57083,  853.91998,   37.00000,   40.20000,   11.70000, 4482.00000
> Building SDMs for SDM algorithms -  CTA','MAXENT.Phillips' , using 10 replicates per algorithm
> Number of background points:  10000> Building final ensemble model with passed thresholds ROC> 0.75 ...
>>> Output SDMs (present and future) saved and plotted
> SDMs completed...

--------------------------------------------------------------
Assessing population structure and imputing missing data...
--------------------------------------------------------------
> Running sNMF population structure analysis...
> Range of K tested: 2 : 10
> K with lowest cross-entropy: 3
>>> Output sNMF population assignment data saved and plotted
> sNMF population structure analysis completed successfully...

> Imputing missing data. using LEA..
> Imputed data saved, LEA completed successfully...

> Imputing missing data using mean genotypes..
> Imputed data saved, mean genotype imputation completed successfully...

> Running PCA population structure analysis...
>>> Output PCA population assignment data saved and plotted

```
** WARNING: check this plot to identify number of population clusters before going further
> PCA  population structure analysis completed successfully...

--------------------------------------------------------------
Running LFMM (Latent Factor Mixed Models)...
--------------------------------------------------------------
> Assessing correlations of predictor variables...
> Correlations of predictor variables checked and completed...
> Variance Inflation Factors for variables are:  2.100223 2.100223
>>> Output correlation data saved and plotted

> Running LFMM (Latent Factor Mixed Models)...
** WARNING: Decision needed for number of populations...

> Evaluate sNMF and PCA population structure outputs to define K
> Predictor variable bioclim_5 ...
> Calculated GIF for predictor variable bioclim_5  is:  1.225879
** WARNING: Examine the unadjusted vs. adjusted p-values, if adjustment is too conservative (distribution is too flat, increase the GIF...)
> Updated GIF is:  0.7723037  (original GIF* 0.63 )
>>> Output summary of adjusted vs. unadjusted p-values plotted
>>> Output Manhattan plot of adjusted p-values saved...
> Predictor variable bioclim_18 ...
> Calculated GIF for predictor variable bioclim_18  is:  1.166093
** WARNING: Examine the unadjusted vs. adjusted p-values, if adjustment is too conservative (distribution is too flat, increase the GIF...)
> Updated GIF is:  0.7346387  (original GIF* 0.63 )
>>> Output summary of adjusted vs. unadjusted p-values plotted
>>> Output Manhattan plot of adjusted p-values saved...
>>> Output candidate SNPs by predictor saved
>>> Output candidate SNPs (merged) saved
> LFMM analysis completed successfully...

--------------------------------------------------------------
Running RDA (Redundancy Analysis)...
--------------------------------------------------------------
** WARNING: Decision needed - how many axes to retain for the RDA? Default is to match the number of predictors - in this case:  2 ( bioclim_5 ,
bioclim_18 )
>>> Output candidate SNPs saved and plotted
> Calculated GIF for predictor variables is:  1.364359
> Updated GIF is:  1.227923  (original GIF* 0.9 )
>>> Output summary of adjusted vs. unadjusted p-values plotted
>>> Output plot of candidate SNPs (post cut-off) saved and plotted
> RDA analysis completed successfully...

--------------------------------------------------------------
Categorising individuals (using RDA)...
--------------------------------------------------------------
Reading in environmental and genomic data...
Subsetting adaptive loci present in both LFMM (FDR<0.05) and RDA (SD<3) analyses...
Number of loci:  26
Performing RDA to categorise individuals...
Writing output plots...
> Categorisation of individuals using RDA completed successfully...

--------------------------------------------------------------
Quantifying adaptive diversity ...
--------------------------------------------------------------
> Reading categorised individuals based on step 9a...
> Summarising samples at each unique lat/long coords ...
> Plotting categorised individuals ...
> DONE ...
> Plotting categorised populations ... (all unique lat/long coords with > 1 sample)...
> DONE ...
> Plotting individual categorisation maps...
> Output plots and data written...

> Adaptive sensitivity analyses completed successfully...
```

#### References

Anantharaman, R., Hall, K., Shah, V., & Edelman, A. (2019). *Circuitscape in Julia: High Performance Connectivity Modelling to Support Conservation Decisions* (arXiv:1906.03542). arXiv. https://doi.org/10.48550/arXiv.1906.03542

Araújo, M. B., Anderson, R. P., Márcia Barbosa, A., Beale, C. M., Dormann, C. F., Early, R., Garcia, R. A., Guisan, A., Maiorano, L., Naimi, B., O'Hara, R. B., Zimmermann, N. E., & Rahbek, C. (2019). Standards for distribution models in biodiversity assessments. *Science Advances*, *5*(1), eaat4858. https://doi.org/10.1126/sciadv.aat4858

Araújo, M. B., & New, M. (2007). Ensemble forecasting of species distributions. *Trends in Ecology & Evolution*, *22*(1), 42–47. https://doi.org/10.1016/j.tree.2006.09.010

Caye, K., Jumentier, B., Lepeule, J., & François, O. (2019). LFMM 2: Fast and Accurate Inference of Gene-Environment Associations in Genome-Wide Studies. *Molecular Biology and Evolution*, *36*(4), 852–860. https://doi.org/10.1093/molbev/msz008

Floyd, C. M. (2020). Introduction to grobblR. Available from: https://cran.r-project.org/web/packages/grobblR/vignettes/grobblR.html.Forester, B. R., Lasky, J. R., Wagner, H. H., & Urban, D. L. (2018). Comparing methods for detecting multilocus adaptation with multivariate genotype–environment associations. *Molecular Ecology*, *27*(9), 2215–2233. https://doi.org/10.1111/mec.14584

Frichot, E., & François, O. (2015). LEA: An R package for landscape and ecological association studies. *Methods in Ecology and Evolution*, *6*(8), 925–929. https://doi.org/10.1111/2041-210X.12382

Frichot, E., Mathieu, F., Trouillon, T., Bouchard, G., & François, O. (2014). Fast and Efficient Estimation of Individual Ancestry Coefficients. *Genetics*, *196*(4), 973–983. https://doi.org/10.1534/genetics.113.160572

Oksanen, J., Blanchet, F. G., Kindt, R., Legendre, P., Minchin, P. R., O'Hara, R. B., Simpson, G. L., Sólymos, P., Stevens, M. H. H., & Wagner, H. (Directors). (2012). *vegan: Community Ecology Package*.

Razgour, O., Forester, B., Taggart, J. B., Bekaert, M., Juste, J., Ibáñez, C., Puechmaille, S. J., Novella-Fernandez, R., Alberdi, A., & Manel, S. (2019). Considering adaptive genetic variation in climate change vulnerability assessment reduces species range loss projections. *Proceedings of the National Academy of Sciences*, *116*(21), 10418–10423. https://doi.org/10.1073/pnas.1820663116

Razgour, O., Taggart, J. B., Manel, S., Juste, J., Ibáñez, C., Rebelo, H., Alberdi, A., Jones, G., & Park, K. (2018). An integrated framework to identify wildlife populations under threat from climate change. *Molecular Ecology Resources*, *18*(1), 18–31. https://doi.org/10.1111/1755-0998.12694

Thuiller, W., Lafourcade, B., Engler, R., & Araújo, M. B. (2009). BIOMOD – a platform for ensemble forecasting of species distributions. *Ecography*, *32*(3), 369–373. https://doi.org/10.1111/j.1600-0587.2008.05742.x

Zizka, A., Silvestro, D., Andermann, T., Azevedo, J., Duarte Ritter, C., Edler, D., Farooq, H., Herdean, A., Ariza, M., Scharn, R., Svantesson, S., Wengström, N., Zizka, V., & Antonelli, A. (2019). CoordinateCleaner: Standardized cleaning of occurrence records from biological collection databases. *Methods in Ecology and Evolution*, *10*(5), 744–751. https://doi.org/10.1111/2041-210X.13152

Zurell, D., Franklin, J., König, C., Bouchet, P. J., Dormann, C. F., Elith, J., Fandos, G., Feng, X., Guillera-Arroita, G., Guisan, A., Lahoz-Monfort, J. J., Leitão, P. J., Park, D. S., Peterson, A. T., Rapacciuolo, G., Schmatz, D. R., Schröder, B., Serra-Diaz, J. M., Thuiller, W., … Merow, C. (2020). A standard protocol for reporting

species distribution models. Ecography, 43(9), 1261–1277.
https://doi.org/10.1111/ecog.04960
